# Diverse Epithelial Lymphocytes in Zebrafish Revealed Using a Novel Scale Biopsy Method

**DOI:** 10.1101/2023.08.09.552511

**Authors:** Gilseung Park, Clay A. Foster, Megan Malone-Perez, Ameera Hasan, Jose Juan Macias, J. Kimble Frazer

**Affiliations:** Depts. of Cell Biology, University of Oklahoma Health Sciences Center, OK, USA; Depts. of Pediatrics, Section of Pediatric Hematology-Oncology, University of Oklahoma Health Sciences Center, OK, USA; Depts. of Microbiology & Immunology, University of Oklahoma Health Sciences Center, OK, USA

## Abstract

Zebrafish are a compelling model to study lymphocytes because zebrafish and humans have similar adaptive immune systems. Human and zebrafish lymphocyte types are conserved, but many aspects of zebrafish lymphocyte biology remain uninvestigated, including lymphocytes in peripheral tissues, like epidermis. Here, we report the first study focused on zebrafish scale epidermal lymphocytes. Zebrafish scales represent a source to longitudinally sample live fish. We developed a novel biopsy technique, collecting scales to analyze epithelial lymphocytes from several fluorescently-labeled lines. We imaged scales via confocal microscopy and demonstrated multiple lymphocyte types in scales/epidermis, quantifying them flow cytometrically. We profiled gene expression of scale, thymic, and marrow lymphocytes from the same animals, revealing B- and T-lineage signatures. Single-cell qRT-PCR and RNA sequencing (scRNAseq) show not only canonical B and T cells, but also novel lymphocyte populations not described previously. To validate longitudinal scale biopsies, we serially sampled scales from fish treated with dexamethasone (DXM), demonstrating epidermal lymphocyte responses. To analyze cells functionally, we employed a bead-ingestion assay, showing thymic, marrow, and epidermal lymphocytes have phagocytic activity. In summary, we establish a non-lethal technique to obtain zebrafish lymphocytes, providing the first quantification, expression profiling, and functional data from epidermal lymphocytes in the zebrafish model.

**Summary:** This study describes a new biopsy method to acquire zebrafish lymphocytes for *ex vivo* studies, without euthanasia. Expression profiles of individual lymphocytes from multiple zebrafish transgenic lines reveal diverse lymphocyte populations, including novel cells expressing genes of both the B- and T-lineages.

## Introduction

Immune cell functions are well-established in skin,^1–5^ which serves as both a barrier and a lymphoid organ.^1,3^ Mammalian cutaneous lymphocytes include epidermal cytotoxic T (CTL) and dermal B, T, and natural killer (NK) cells.^6–9^ Zebrafish share B/T/NK lymphocytes, surface immunoglobulin (sIg), T cell receptors (TCR), MHC molecules, and other immunologic features with mammals.^10–12^ Zebrafish lymphocytes are well-characterized,^13–20^ but most studies focus only on cells of lymphoid tissues. Several reports briefly mention epidermal lymphocytes: one noted *igm^+^* skin B cells;^19^ another *mpeg1.1*^+^ B cells;^14^ a third identified B-1-like cells in multiple organs, including skin.^21^ However, these reports concentrate on thymus, marrow, and spleen, while most skin/scale studies investigate osteogenesis^22,23^ or melanoma.^24^

Zebrafish are small, making non-lethal lymphocyte collection difficult. Obtaining blood retro-orbitally^25^ is technically-challenging, with serial-sampling infeasible. Instead, most reports (including ours^18,26,27^) collect lymphocytes post-euthanasia, precluding longitudinal studies. Here, we describe a new method to collect 1-20 scales/fish, yielding hundreds-to-thousands of lymphocytes and permitting repeated sampling. Via scale biopsy, we examined several transgenic lines, identifying, quantifying, and expression-profiling epidermal lymphocytes. Single-cell studies revealed B- and T-lineage subtypes and potentially novel lymphocyte populations. We also functionally tested lymphocyte DXM-responses and phagocytosis. Overall, we describe a simple technique to collect zebrafish epidermal lymphocytes and their first comprehensive study.

## Materials & Methods

### Zebrafish

Zebrafish care was reported previously,^18,26,27^ with a 14:10-hour circadian-cycled colony kept at 28.5°C. Experiments followed IACUC-approved protocols. Anesthesia utilized 0.02% tricaine methanesulfonate (MS-222). Transgenic *cd79:GFP*^15^ and *lck:mCherry* fish (from the Zon laboratory, Harvard University) were inter-bred, creating double-transgenics.

### Biopsies

Scale biopsy is detailed in Supplemental Materials.

### Microscopy

Anesthetized 3-to-9m fish were imaged with a Nikon AZ100 microscope. Low- and high-exposure images were obtained as previously^18,26^ using a Nikon DS-Qi1MC camera and NIS Elements (V4.13) software. Select images are brightened to enhance dim-fluorescence.

### Confocal imaging

Scales were slide-mounted with SlowFade™ Glass Soft-set Antifade Mountant plus DAPI. Imaging utilized a Leica-SP8 confocal microscope and LAS-X (V3.7.4.23463) software.

### Flow cytometry/FACS

Tissues were pestle-dissociated and passed through 35μm filters.^26^ Cells were collected from lymphoid/precursor gates using a BD-FACSJazz (Becton-Dickinson). Flow cytometry used a CytoFLEX^TM^ and Kaluza software (Beckman-Coulter).

### qRT-PCR

RNA was extracted using Trizol (Invitrogen). Total RNA triplicates (16ng) were reverse-transcribed for SYBR-Green qRT-PCR with a CFX96 Touch^TM^-PCR System (Biorad). Two^-ΔCt^ (Ct_Δ_ = Ct_experimental_−Ct_housekeeping_)^28^ calculated expression relative to *β-actin* and *eef1a1l1*.

### Single-cell qRT-PCR

Cells were FAC-sorted into 96-well plates, lysed, and cDNA synthesized using Fluidigm™ protocols. Supplementary Material lists sc-qRT-PCR details. CT values were converted to Log2Ex using a limit of detection (LoD) = 27. Other LoDs yielded similar results. Cells expressing >30% T- and <30% B-lineage genes were designated T-lineage, and vice versa. Cells expressing >30% of T- and B-lineage genes were designated Bi-Phenotypic.

### scRNAseq

Cell-capture utilized 10xGenomics Chromium, with Illumina NovaSeq6000 sequencing. Cells (18,000 thymic; 8,800 marrow; 3,500 scale) were loaded (1,000 cells/µl) on 10xGenomics single-cell v2 chips into a Chromium controller and processed per 10xGenomics protocol. Supplementary Material details data-processing and analysis steps.

### DXM treatment

Zebrafish were continuously-housed for 10d in 10μg/ml DXM in 500ml fish water, changed daily, as previously.^27^ Fish were imaged pre- and post-treatment, with fluorescence-integrated densities quantified via ImageJ.^29^

### Phagocytosis assays

Fluorescent lymphocytes were sorted into 1.5ml Eppendorfs in 5μl volumes with a BD-FACSAria Fusion (Becton-Dickinson), incubated with 1μl of 1:1000-diluted 1μm blue-FluoSpheres (Thermo-Fisher) for 30min, and analyzed via ImageStream™ MkII and IDEAS software (Luminex).

### Data Sharing Statement

scRNA-seq data used in this study are available at GEO under accession number GSE237417.

## Results

### Zebrafish epidermal lymphocytes

To investigate zebrafish epidermal lymphocytes, we analyzed transgenic fish bearing lymphocyte-specific fluorophores. Microscopy of B-labeled lines, *cd79a:GFP* and *cd79b:GFP*,^15^ exhibited head and gill fluorescence; T cell-specific *lck:GFP* fish displayed thymic fluorescence (Figure 1A), as previously described.^13,26,27^ To determine whether fluorescence emanated from skin/scales, we developed a novel scale biopsy method (details in Supplemental Material).

**Figure 1:**
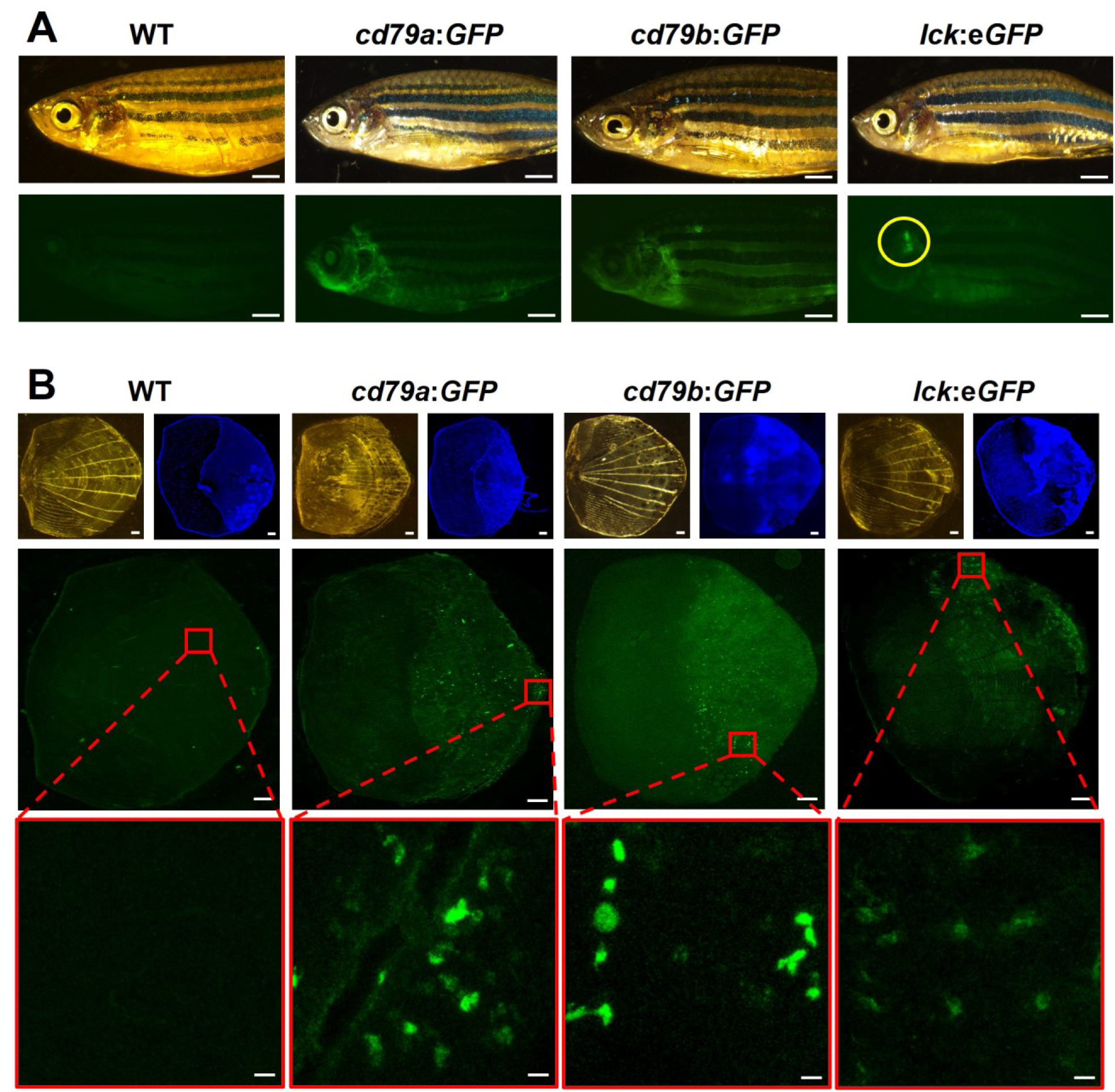
Zebrafish lymphocytes in scale epidermis. **(A)** Light and fluorescent microscopy images of wild-type (WT) or transgenic *cd79a:GFP*, *cd79b:GFP*, and *lck:GFP* fish. Yellow circle denotes thymic region. White bars = 2mm. **(B)** Top row: light microscopy images of single scales from each fish in panel 1A. DAPI highlights nuclei concentrated in caudal, water-exposed region of scales. Middle row: low-power confocal fluorescent images show GFP^+^ cells in the epidermis of all three transgenic lines. Bottom row: red-boxed regions in middle rows at high-power. White bars = 100 μm in light microscopy, DAPI, and low-power confocal images or 10 μm in high-power images.

Upon removal, epidermis remains attached to lateral/exterior and medial/interior scale surfaces (Figure 1B, DAPI).^22^ Expectedly, by fluorescent microscopy, wild-type (WT) scales lacked fluorescence (Figure 1B, left) except rare auto-fluorescent cells (Figure S1A). However, *cd79a:GFP*, *cd79b:GFP*, and *lck:GFP* epidermis (Figure 1B, right three columns) were all fluorescent. Confocal imaging revealed GFP^+^, ∼10μm lymphocytes, some with cytoplasmic protrusions (Figures 1B red-boxes, S1A yellow-boxes), confirming these findings. 3-Dimensional confocal microscopy movies of scales display fluorescent cells from various perspectives (Figure S1B).

### Identifying epidermal lymphocytes

We next analyzed fluorescent cells from scales of these and two additional transgenic lines: dual-transgenic *lck:GFP*;*rag2:hMYC* fish have lymphocytosis and eventually develop acute lymphoblastic leukemias;^18,26,27^ we also tested *lck:mCherry* fish (gift from the Zon laboratory, Harvard University). We identified lymphoid gates^30^ by forward- and side-scatter (Figure 2A) and detected fluorescent epidermal cells from each line (Figure 2B). B-labeled lines (*cd79a:GFP*, *cd79b:GFP*) exhibited GFP^lo^ and GFP^hi^ populations, (Figure 2B, upper-histograms). T-labeled *lck:GFP* (+/- *rag2:hMYC*) and *lck:mCherry* scales had only GFP^lo^/mCherry^lo^ cells (Figure 2B, lower-histograms). Transgenic human *MYC* (*hMYC*) did not alter GFP intensity, but slightly increased GFP^lo^ cells/scale (quantified subsequently).

**Figure 2:**
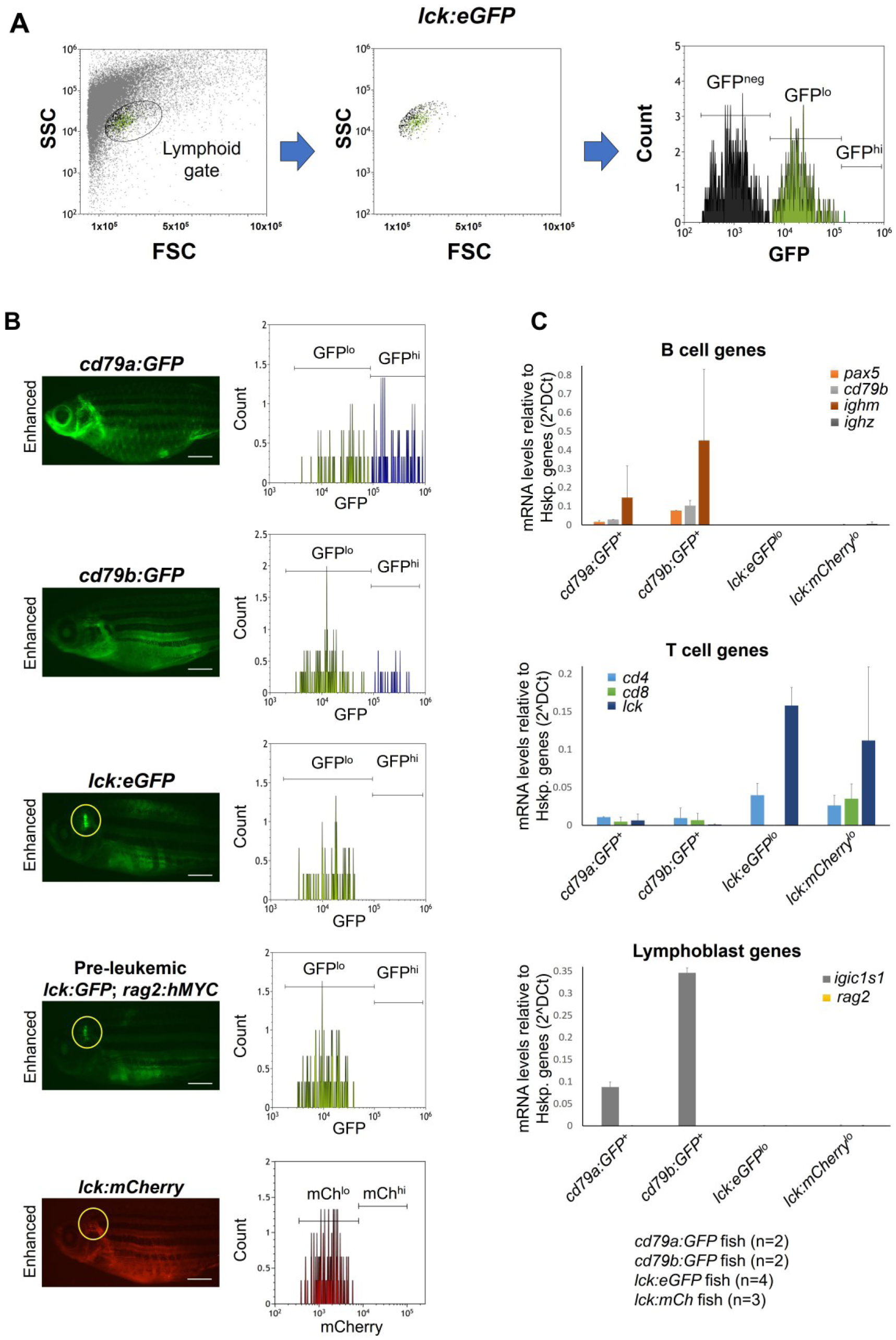
Epidermal lymphocyte identities. **(A)** Flow cytometry work-flow showing example lymphoid and GFP^-^, GFP^lo^, and GFP^hi^ gates in *lck:GFP* fish. **(B)** Fluorescent microscopy images and flow cytometry histograms of scale cells from B- (*cd79a:GFP*, *cd79b:GFP*) or T- (*lck:GFP*, *lck:GFP* + *rag2:hMYC*, *lck:mCherry*) labeled lines. Yellow ovals denote thymic region. White bars = 2mm. (**C**) Bulk qRT-PCR of FAC-sorted GFP^+^ cells from *cd79a:GFP* (n=2), *cd79b:GFP* (n=2), *lck:GFP* (n=4), or *lck:mCherry* (n=3) fish (40 scales/fish) using B- (*pax5*, *cd79b*, *ighm*, *ighz*), T- (*cd4*, *cd8*, *lck*), or lymphoblast- (*igic1s1*, *rag2*) specific genes. Results normalized to housekeeping genes *eef1a1l1 (ef1a)* and *rpl13a*, shown as mean ± S.D.

To confirm cell identities, we measured B- (*pax5, cd79b, ighm*, *ighz*), T- (*cd4, cd8, lck*), and lymphoblast- (*igic1s1*, *rag2*) transcripts by qRT-PCR (primers in Supplementary Table 1). GFP^+^ cells (GFP^lo^ + GFP^hi^) of both *cd79* lines expressed B-lineage mRNAs, unlike GFP^+^/mCherry^+^ cells of *lck* fish (Figure 2C, top). Thus, both *cd79* lines label the B-lineage, further proving that B cells reside in zebrafish epidermis. Oppositely, *lck* lines’ fluorescent cells expressed T-lineage genes, unlike both *cd79* lines (Figure 2C, middle), demonstrating T cells also populate scales. Both *cd4* and *cd8* were detected in *lck*:*mCherry* scales, indicating both T_H_ and CTL are present. However, *cd8* was not detected in *lck*:*GFP* scale cells, suggesting a difference between the *lck* lines. Scale lymphocytes lacked *rag2* (Figure 2C, bottom), implying they are mature B and T cells. However, we detected *igic1s1* (which we previously postulated to be a pre-B surrogate Ig light chain^31^) in epidermal B cells of both *cd79* lines. This may reflect residual mRNA in mature B cells, or that our prediction is erroneous. Regardless, these data demonstrate T_H_, CTL, and B cells each reside in zebrafish epidermis.

### Single-cell profiles reveal multiple epidermal lymphocyte types

We next analyzed GFP^hi^ and GFP^lo^ cells from *cd79a* and *cd79b* scales, thymus, and marrow of the same fish via single-cell multiplex-qPCR, using orthologues that distinguish B, T, and other mammalian leukocytes (Figure 3A).^26,27,32^ We tested 16 B- (*pax5*, *cd79a*, etc.) and 10 T-lineage (*cd4*, *lat*, etc.) transcripts, 17 lymphoblast (*rag1*, *rag2*), leukocyte (*l-plastin*), erythroid (*ba1*), or lymphocyte (*bcl6a*, *id3*, *ets1*) mRNAs, plus housekeeping (*eef1a1l1/ef1a*) and control (*GFP*) genes (primers in Supplementary Table 2).

**Figure 3:**
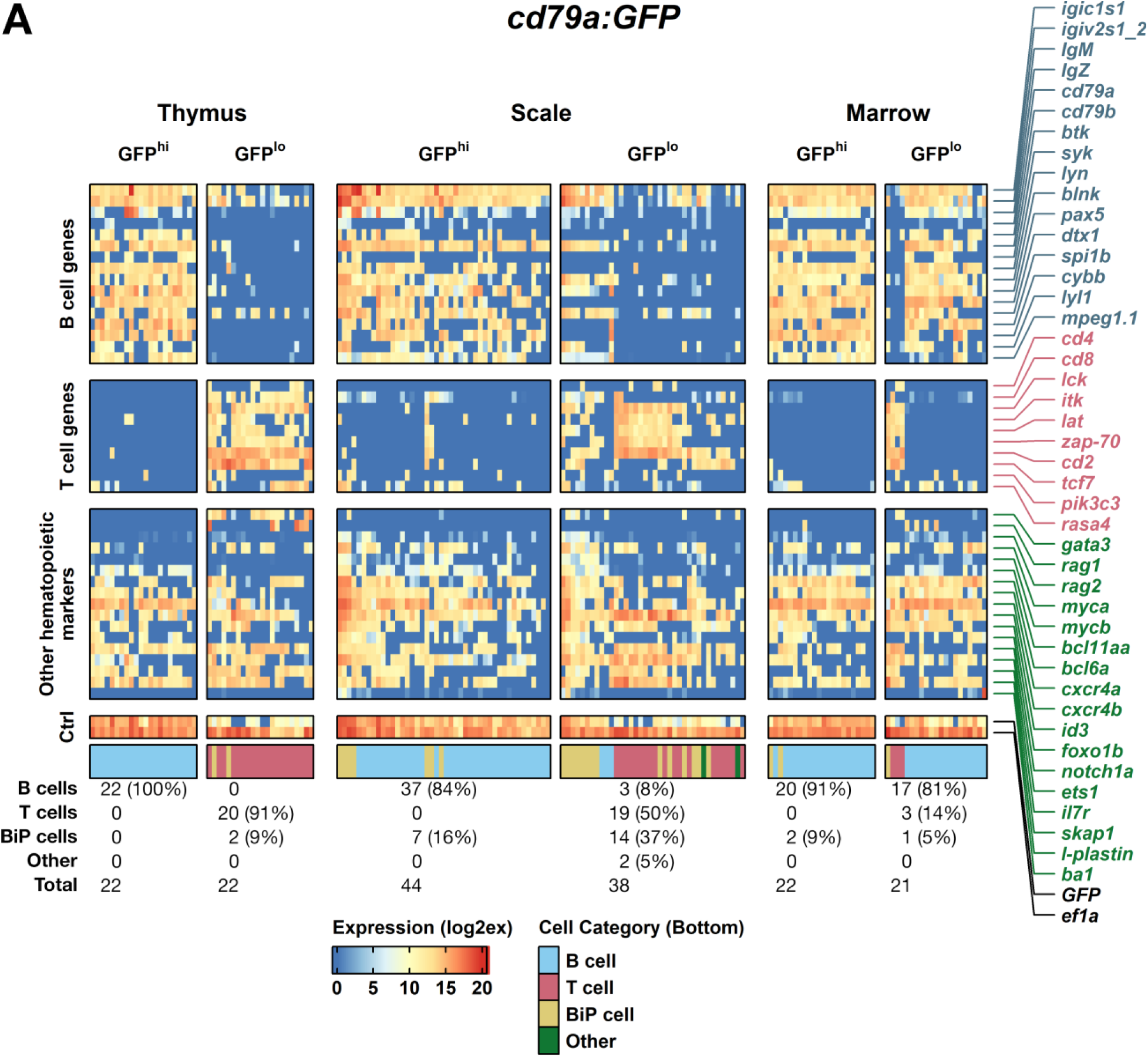
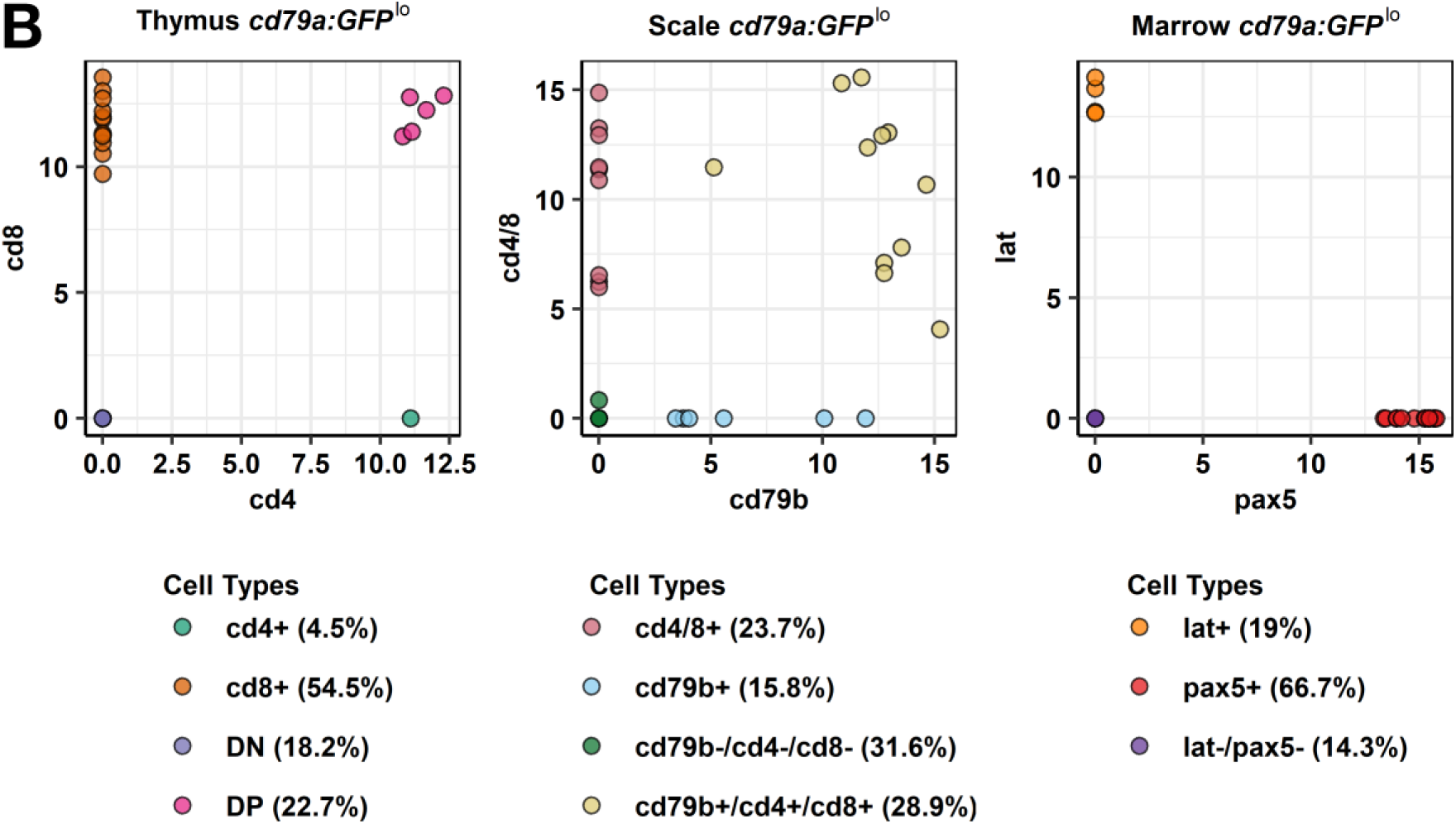
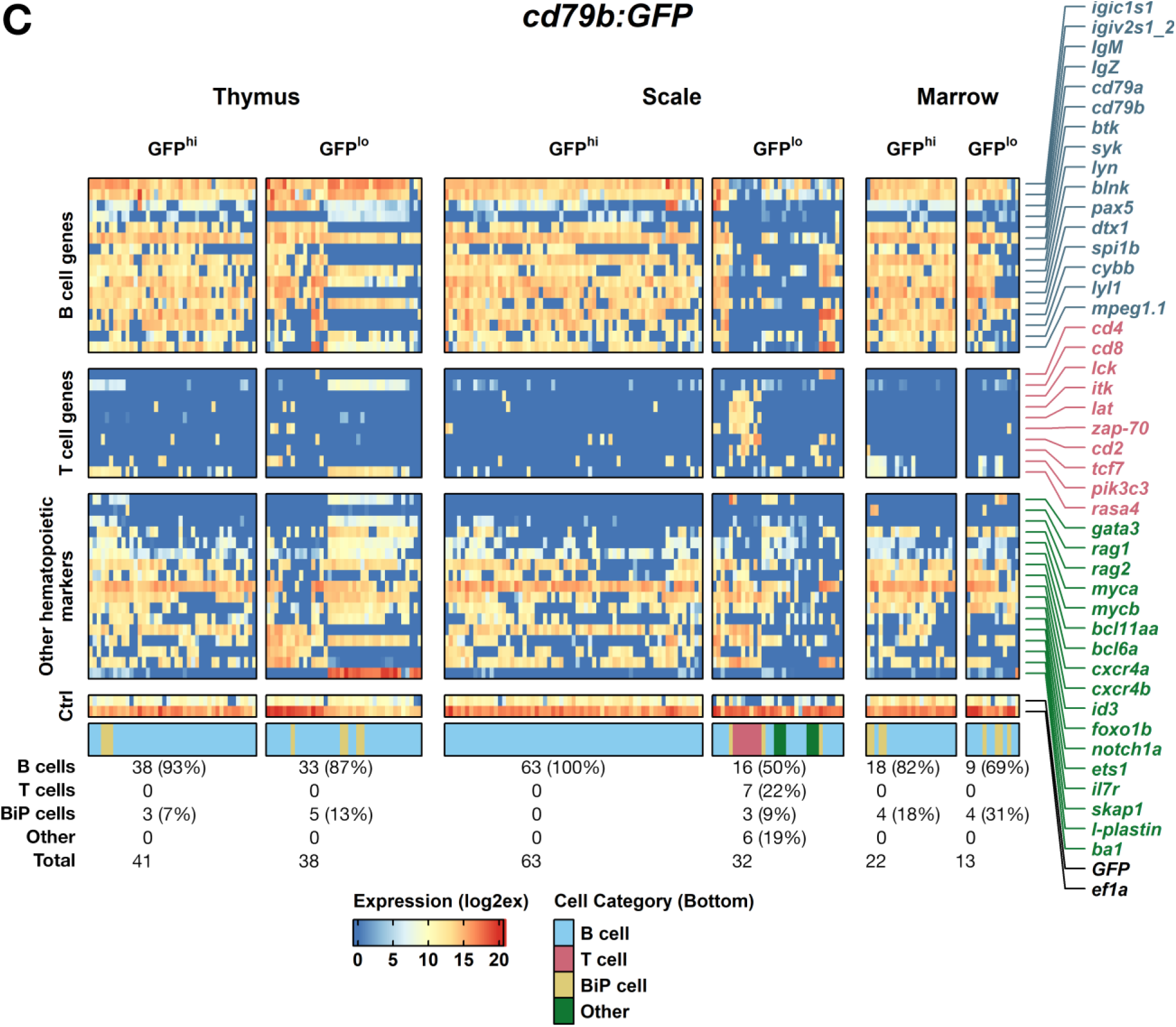
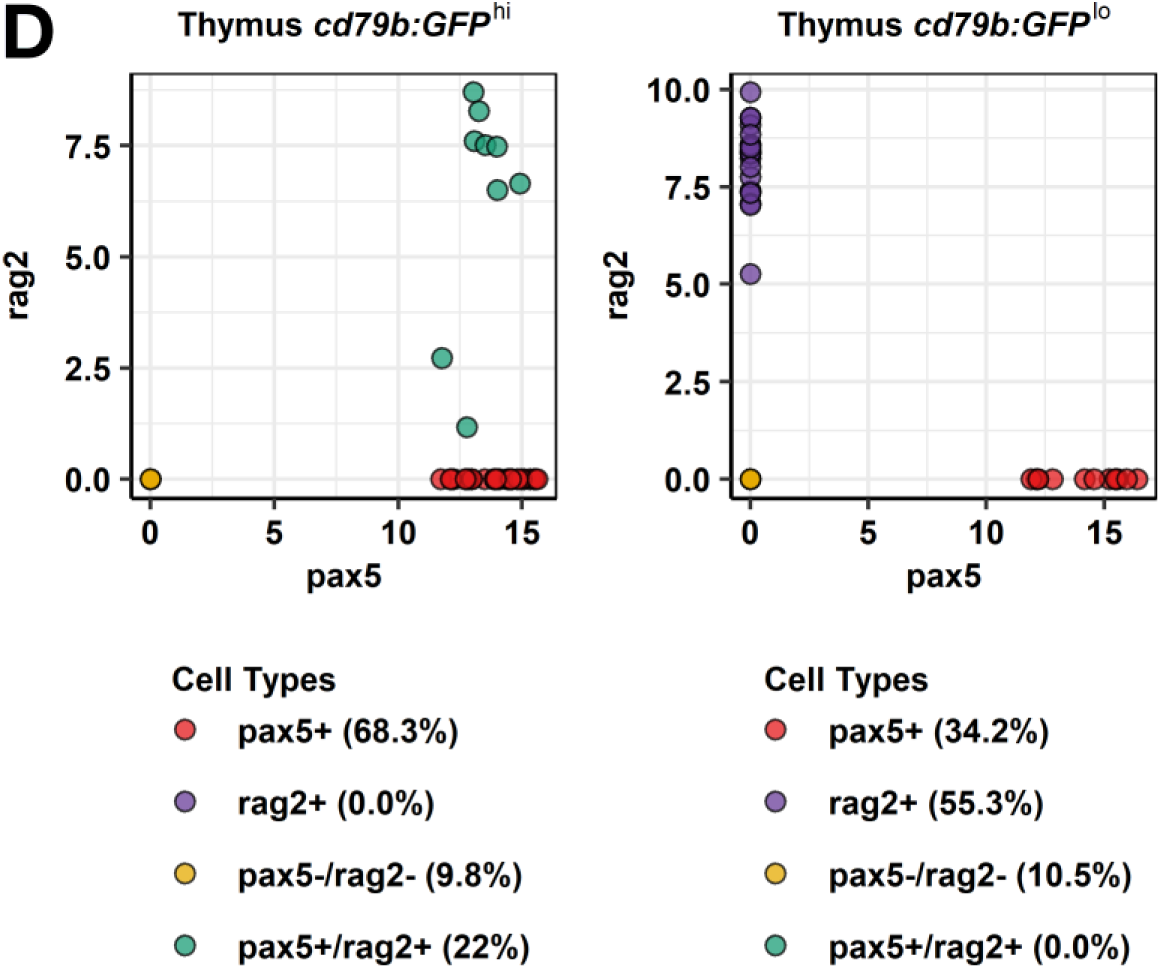
Single-cell gene expression profiles of lymphocytes from *cd79*-labeled fish. **(A)** Transcript levels in GFP^hi^ and GFP^lo^ populations from *cd79a:GFP* thymic, scale, and marrow cells obtained using multiplex-qPCR. Labels at top list genotype, tissue, and GFP intensity of each sample type. Inferred cell identities are below heatmaps with total cell number and percentage. Cell identifies are color-coded in legend below figure. **(B)** Dot-plots depicting expression levels of genes listed on X- and Y-axes. Each dot is one cell from the heatmap in **A**. **(C-D)** Identical data depictions as in **A** and **B**, with results for *cd79b:GFP* fish. Expression of *eef1a1l1* (*ef1a*; bottom heatmap row) was used as a threshold to confirm wells contained amplifiable RNA. CT values converted to Log_2_Ex for visualization of relative expression levels.

GFP^hi^ thymocytes of *cd79a* fish expressed B-lineage genes (*pax5*, *cd79a, cd79b*, *ighm*, etc.), supporting bulk qRT-PCR results, and other B cell receptor (BCR)-related genes *syk*, *lyn*, and *blnk*, but not T-lineage genes. Cells lacked *rag1*/*2*, but 10/22 expressed *il7r*, like pre-B cells.^33^ We designated these GFP^hi^ thymocytes “*pax5*^+^ mature B cells” (Figure 3A, 1^st^-column).

Unexpectedly, *cd79a* GFP^lo^ thymocytes had near-opposite profiles, with mostly T- rather than B-lineage transcripts (Figure 3A, 2^nd^-column). Thus, unexpectedly, they are T cells. Curiously, although GFP^lo^ in the *cd79a:GFP* background, these cells lacked *cd79a* mRNA and had low-to-undetectable *GFP* transcripts. We postulate these cells transcribed *cd79a* and *GFP* previously, but when FACS-purified, only scant GFP protein remains, explaining their GFP^lo^ status. Notably, human T-lymphoblast cancers are occasionally CD79A^+^,^34–36^ suggesting some T-lymphoblasts may express *CD79A*. Alternatively, *cd79a*:*GFP* may imperfectly recapitulate genuine *cd79a* expression. Regarding *cd4* and *cd8*, most GFP^lo^ cells were immature *cd4*/*cd8* double-positive (DP) or mature *cd8* single-positive (SP), with only one *cd4* SP cell (Figure 3B, left).

We simultaneously analyzed scale lymphocytes. Scale GFP^hi^ cells were near-identical to GFP^hi^ B-thymocytes (Figure 3A, 3^rd^ vs. 1^st^-columns), except ∼32% (14/44) had detectable *rag2*, suggesting recent Ig-recombination. Thus, some skin B cells are likely newly-immigrated. Unlike GFP^lo^ thymocytes, which were mostly T-lineage (Figure 3A, 2^nd^-column), GFP^lo^ scale cells (4^th^-column) comprised several cell types: 50% (19/38) resembled GFP^lo^ thymocytes, but only SP (*cd4*/T_H_ or *cd8*/CTL, not DP). Another ∼37% (14/38) were highly unusual, with high levels of both B- (*cd79b*, *lyn*, *blnk*, etc.) and T- (*cd8*, *lck*, *lat*, etc.) lineage mRNAs. We designated these, which express both BCR- and TCR-signaling pathway genes, as “Bi-Phenotypic” (BiP) cells (tan in Figure 3A). Many also expressed *ighm* and/or *ighz*, and *mpeg1.1*, another zebrafish B-lineage marker.^19^ However, BiP cells lacked other B-lineage genes (*pax5*, *btk*, *syk*, *cd79a*) that GFP^hi^ scale/thymic B cells expressed, also dissimilar from previously-described BiP-ALL.^37,38^ To further delineate BiP cells, we ranked GFP^lo^ scale cells using T (*cd4*/*cd8*) versus B (*cd79b*) transcripts (Figure 3B, middle), resulting in ∼29% BiP (tan), ∼24% T (red), ∼16% B (light blue), with ∼32% uncategorized (green). Plotting *cd2* vs. *cd79b* yielded similar results (Figure S2A, upper plot). BiP lymphocytes, which have not been reported previously, were also detected in other populations (tan hashes in Figure 3A, 2^nd^-6^th^ columns, and subsequent figures).

We also analyzed marrow lymphocytes. Like thymus and skin, marrow GFP^hi^ cells were mostly B-lineage (Figure 3A, 5^th^-column), with little *rag1* (1/22) or *rag2* (5/22), suggesting mature B cells transcribe more *cd79a* (thus, they are GFP^hi^). Marrow GFP^lo^ cells differed from GFP^lo^ thymic/scale cells. In marrow, few GFP^lo^ lymphocytes (∼14%) were T-lineage (Figure 3A, 6^th^-column); most cells (∼81%) resembled other tissues’ GFP^hi^ B cells. Marrow GFP^lo^ vs. GFP^hi^ B cell maturity also differed, with just ∼23% of GFP^hi^ B cells *rag1^+^/2^+^* versus ∼48% of GFP^lo^ B cells (again showing B-precursors have lower *cd79a* and GFP). A two-gene index (*lat* for T, *pax5* for B) estimated marrow GFP^lo^ cells at 19% T, 67% B, with 3 unclassified cells (Figure 3B, right). Plotting *cd2* vs. *pax5* was identical (Figure S2A, lower dot plot).

In summary, all *cd79a* GFP^hi^ lymphocytes exhibited unambiguous B-lineage profiles, but GFP^lo^ lymphocytes varied by tissue: GFP^lo^ thymocytes were DP or SP (mostly *cd8*^+^/CTL) T cells. Marrow GFP^lo^ cells were ∼81% B-lineage (∼48% immature *rag*^+^) and ∼14% T-lineage. Scale GFP^lo^ cells were most complex, comprising *cd4*^+^ or *cd8*^+^ SP T cells and mixed-lineage BiP lymphocytes. Thymus and marrow also contained occasional BiP cells. To independently re-test these findings, we next analyzed *cd79b*:*GFP* fish.

GFP^hi^ thymocytes of *cd79b* fish resembled *cd79a* thymic *pax5*^+^ B cells (1^st^-columns of Figure 3C vs. 3A), but dissimilarly, ∼24% were *rag1*^+^ and/or rag*2*^+^. Two-gene analysis (*rag2*, *pax5*) delineated these B-subtypes (Figure 3D, left). Unlike *cd79a* GFP^lo^ thymocytes that were T-lineage, *cd79b* GFP^lo^ thymocytes harbored two B-populations: immature (*pax5*^-^/*rag2*^+^) and mature (*pax5*^+^/*rag2*^-^; Figure 3C, 2^nd^-column), which were mutually-exclusive by two-gene (*rag2*/*pax5*) analysis (Figure 3D, right). GFP^lo^ mature-*pax5*^+^ cells expressed no *ighz* and much higher *ighm* than GFP^lo^ immature-*pax5*^-^ B-thymocytes, which transcribed *ighm* and *ighz* similarly. The immature *pax5*^-^/*rag2^+^* population was clearly B-lineage, highly expressing Ig-variable genes (*igic1s1*, *igiv2s1_2*), and Ig-constant (*ighm*, *ighz*), *cd79b*, *lyn*, *blnk*, and *mpeg1.1* transcripts. However, they also resembled *cd79a* BiP cells that were *pax5*^-^/*cd79a*^-^ and expressed *ighm* and/or *ighz* and *cd8.* Finally, like GFP^hi^ B-precursors, they exhibited high *rag2*, *rasa4*, *gata3*, *il7r* and *myc* orthologues. We speculate these cells are nearing B-lineage commitment, silencing non-B genes and initiating Ig-rearrangement.

GFP^hi^ *cd79b* scale cells (Figure 3C, 3^rd^-column) matched GFP^hi^ *cd79a* scale B cells (Figure 3A, 3^rd^-column) and GFP^hi^ *cd79b* thymocytes (Figure 3C, 1^st^-column), but as in *cd79a* fish, GFP^lo^ scale cells (Figure 3C, 4^th^-column) had multiple populations: (1) mature-*pax5*^+^ B cells (identical to thymic GFP^lo^ *pax5^+^* B cells), (2) *pax5*^-^ B cells (different from thymic *pax5*^-^ B cells, with higher *cd4*, *mpeg1.1*, absent *ba1*, and low-to-absent *ighm*, *ighz*, and other Ig mRNAs), (3) T cells (resembling *cd79a* GFP^lo^ scale cells), and (4) BiP cells resembling *cd79a* GFP^lo^ scale BiP lymphocytes. A fifth unclassifiable group (∼19%) co-expressed low B-/T-lineage gene levels.

Marrow *cd79b* cells confirmed prior results, with >90% GFP^hi^ and >50% GFP^lo^ cells having mature-*pax5*^+^ B-profiles like those seen in thymus/scales (Figure 3C, 5^th^-6^th^ columns). Remaining GFP^lo^ marrow cells were *pax5*^-^, yet expressed other B-lineage genes. Unlike *cd79a*, no marrow GFP^lo^ cells were T-lineage in *cd79b* fish. Overall, most GFP^hi^ cells (198/214; ∼93%) in every tissue from both *cd79*-labeled lines were B-lineage; B-subtypes varied by tissue and genotype. Another ∼7% (16/214) showed BiP features. Both lines’ GFP^lo^ cells were more complex, containing B, T, and BiP cells; subtypes (e.g., *pax5*^+^ vs. *pax5*^-^) also varied by tissue and genotype. We also analyzed GFP^-^ cells. Many were not convincingly lymphoid, although some GFP^-^ B, T, and BiP cells were detected (Figure S2B).

### Single-cell profiling double-transgenic fish

We next bred *cd79a*:*GFP* to *lck*:*mCherry* fish to make double-transgenics that resembled both parental lines (compare Figures 4A and 1A). From these, we purified four populations: P1—GFP^hi^/mCh^-^; P2—GFP^lo^/mCh^hi^; P3—GFP^-^/mCh^hi^; P4—GFP^-^/mCh^lo^; abundance varied by tissue (Figure 4A, Table 1). sc-qPCR revealed multiple profiles (Figure 4B). P1 was exclusively B-lineage in every tissue. P2 resembled *cd79a* GFP^lo^ thymocytes, with two differences: (1) DP cells were absent (versus 23% of *cd79a* GFP^lo^ thymocytes; Figure 3B vs. S3A) with most being SP *cd8*^+^/CTL. (2) Few P2 cells were *gata3*^+^, versus 68% of *cd79a* GFP^lo^ thymocytes. Mammalian T_H_2, invariant NK/T2 (iNKT2), and innate lymphoid cell-2 (ILC2) cells express GATA3, perhaps explaining this.^39,40^

**Figure 4:**
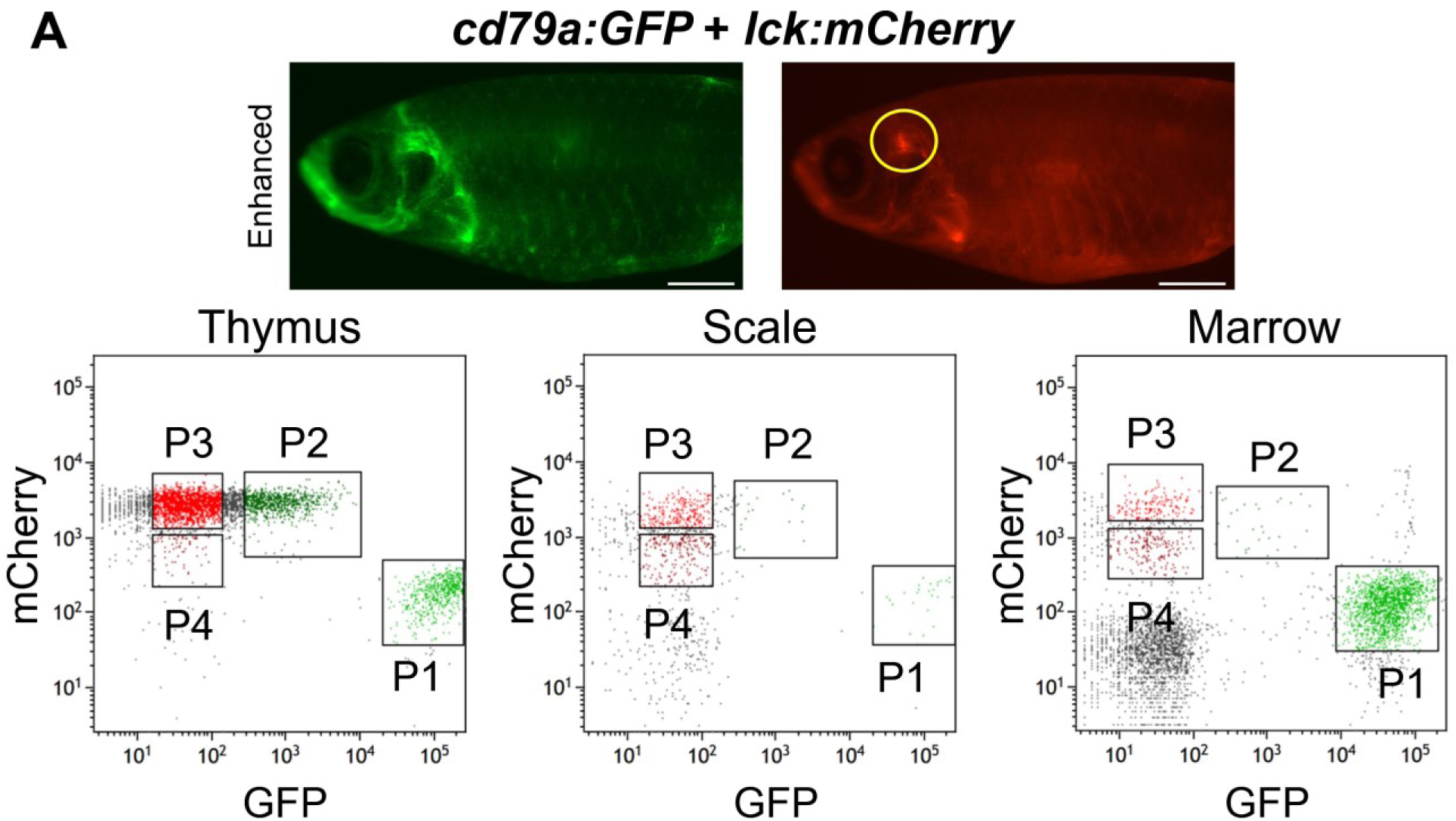
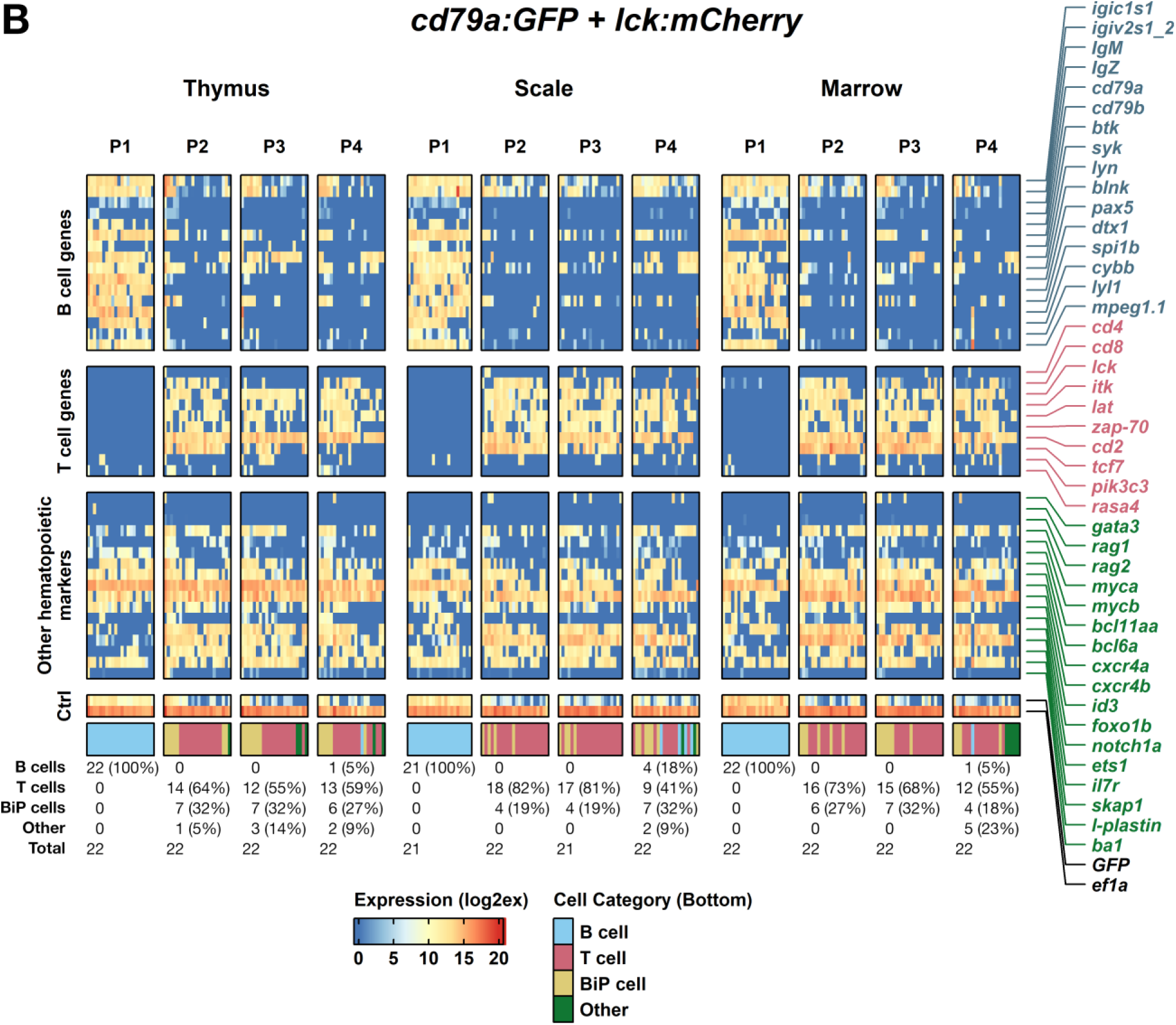
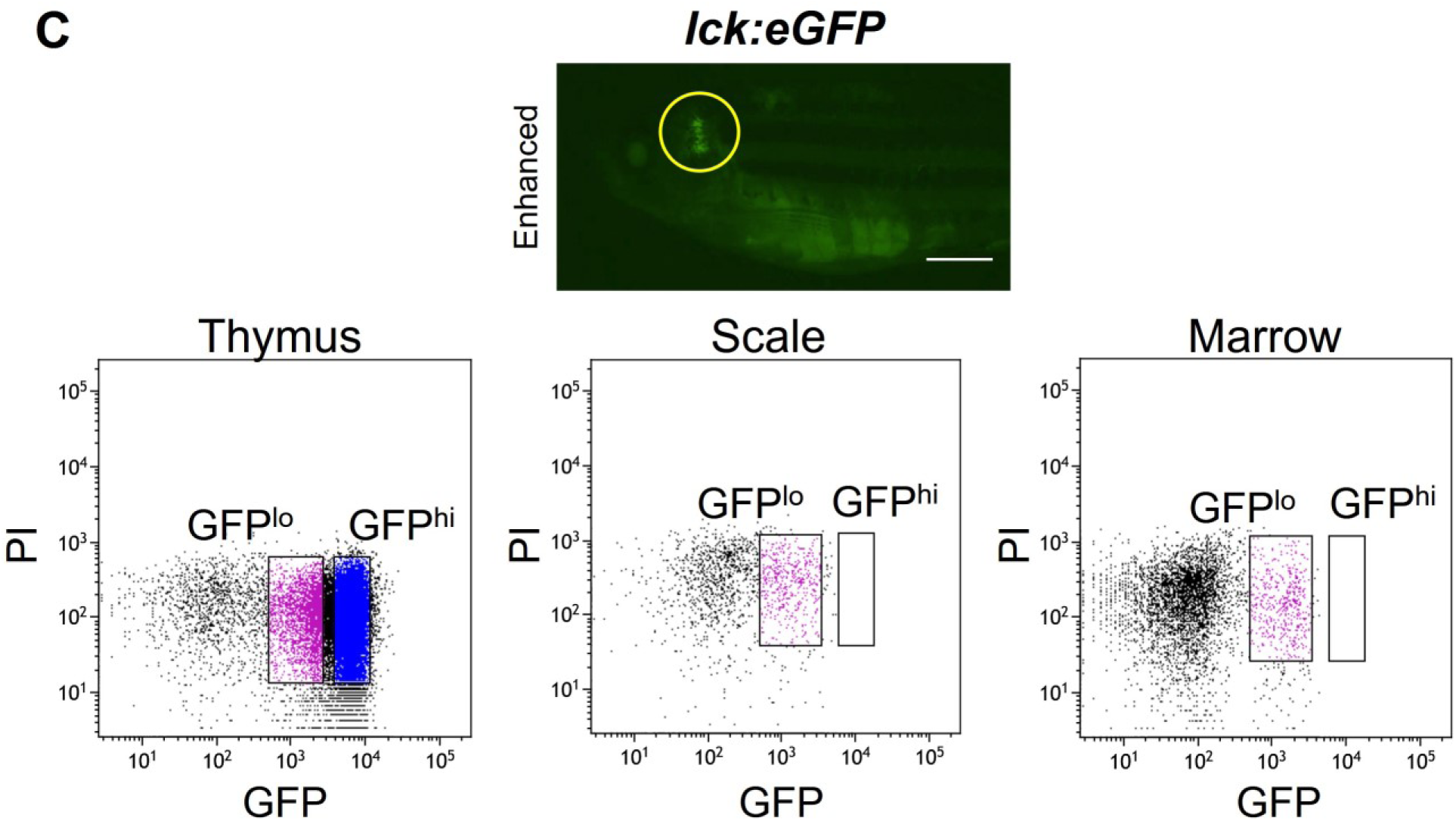
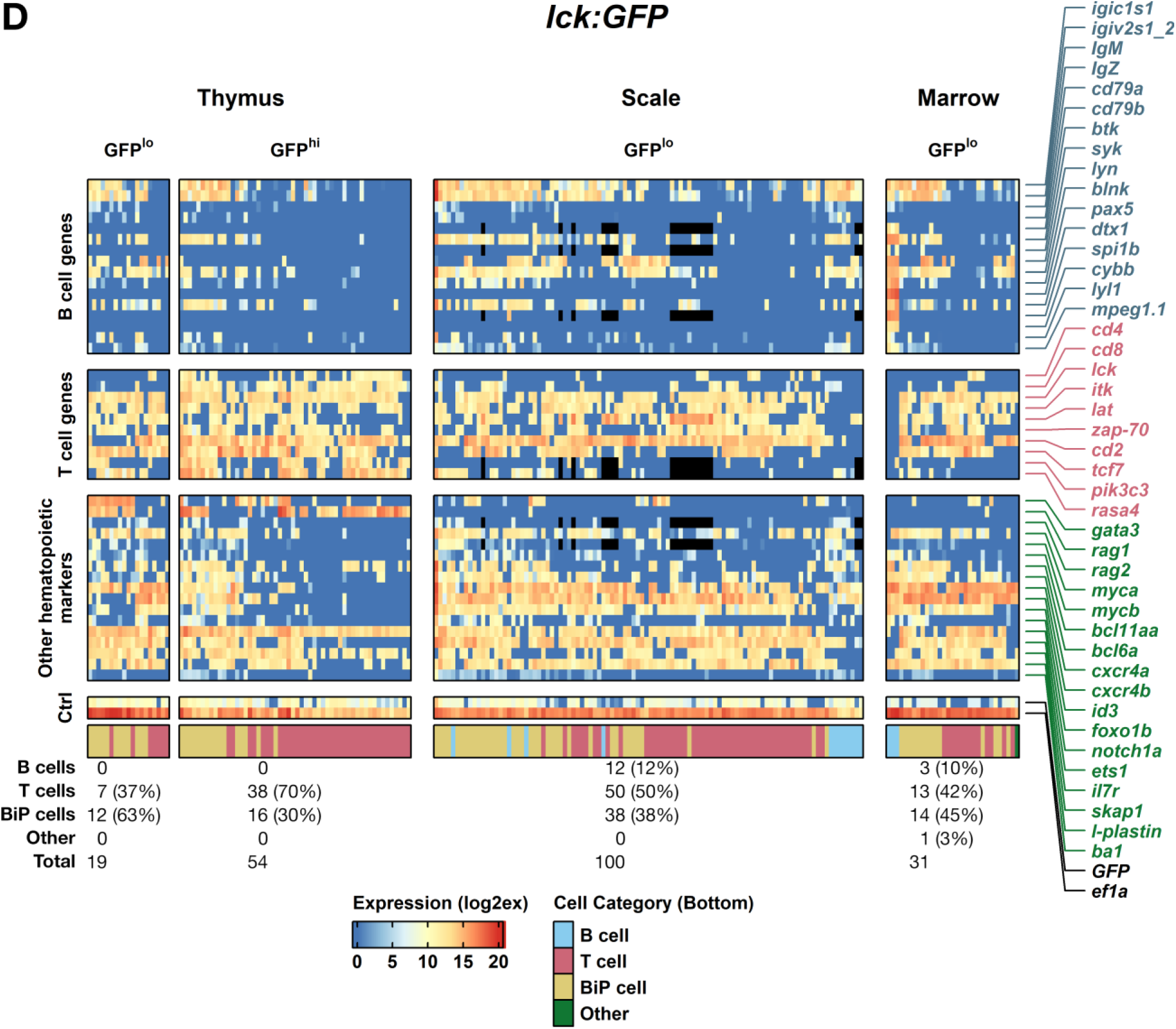
Single-cell profiles of lymphocytes from *cd79a:GFP* + *lck:mCherry* and *lck:GFP* fish. **(A)** Fluorescent microscopy images and flow plots of thymus, scale, and marrow from *cd79a:GFP* + *lck:mCherry* fish. Yellow oval denotes thymic region. White bars = 2mm. Populations 1-4 are described in the main text. **(B)** Transcript levels in Populations 1-4 from each tissue (left, thymus; middle, scale; right, marrow). Data depiction identical to Figures 3A and **3C**. **(C)** Same format as in **A**, with results for *lck:GFP* fish. Propidium iodide (PI; Y-axis) used to verify viability. **(D)** Data depiction identical to **B**, with transcript levels for GFP^lo^ and GFP^hi^ cells. Black boxes in center (scale) column indicate genes not tested.

**Table 1:**
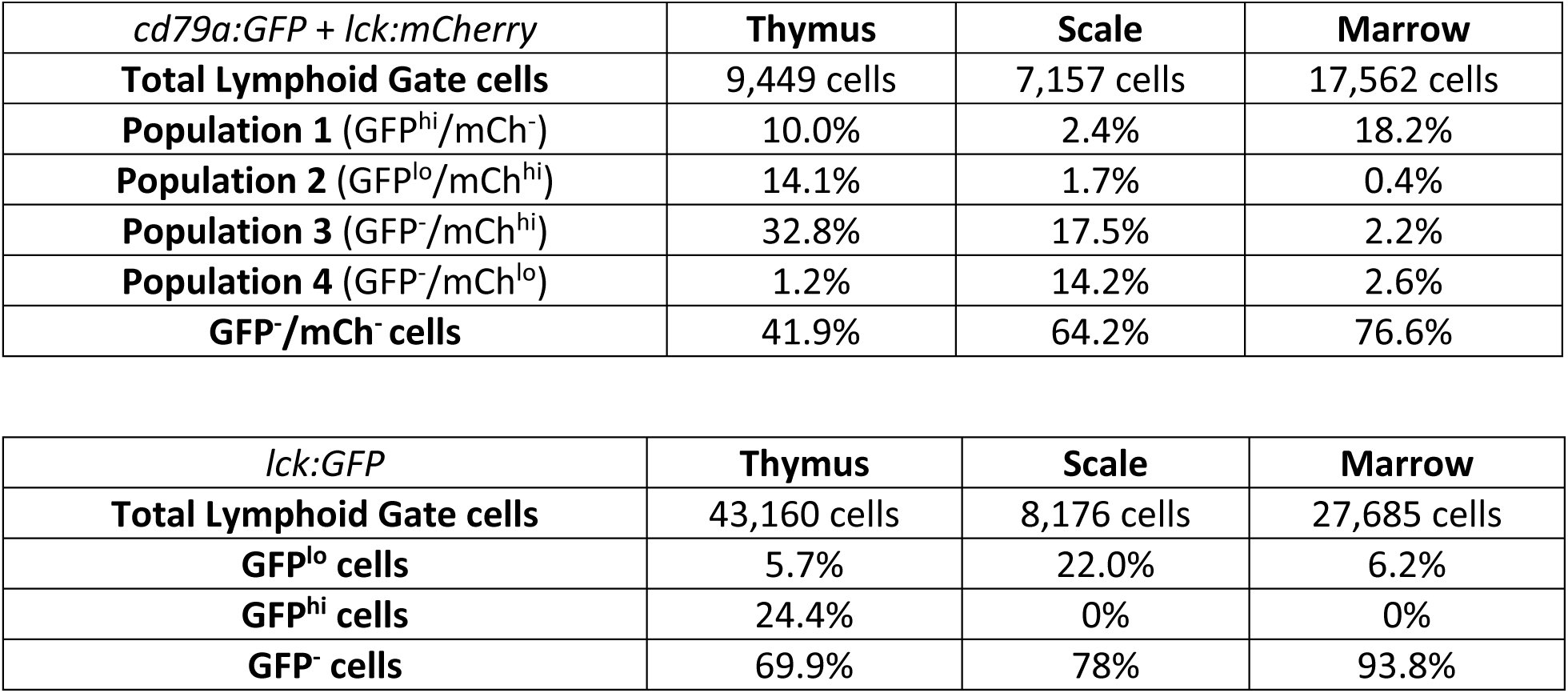
Cell population frequencies in *cd79a:GFP* + *lck:mCherry* and *lck:GFP* zebrafish.

GFP^-^ thymocytes that differed by mCherry level (P3, mCh^hi^; P4, mCh^lo^) were both primarily T-lineage, resembling P2. Many P2-P4 thymocytes also expressed B-lineage transcripts (*cd79b*, *syk*, *lyn*, *blnk*), like BiP cells (Figure 4B, thymus P2-P4 columns. Scale and marrow P2-P4 mimicked thymic P2-P4, with many cells also having BiP-profiles (Figure 4B, scale/marrow P2-P4 columns).

We also analyzed *lck:GFP* fish, whose thymic fluorescence resembled *lck:mCherry* (Figure 4A, 4C). Thymi had large GFP^hi^ and smaller GFP^lo^ fractions, but unlike mCherry scales/marrow that contained both mCh^hi^ and mCh^lo^ cells (Figure 4A), *lck:GFP* scales/marrow contained no GFP^hi^ cells (Figure 4C). By sc-qPCR, *lck:GFP* GFP^lo^ thymocytes had strong T-signatures, like P4 (mCh^lo^) thymocytes (Figure 4D 1^st^-column vs. 4B 4^th^-column). However, ∼80% were *rag1*^+^/*2*^+^, versus no thymic P4 cells. These *rag*^+^ cells had BiP features, expressing Ig (*igic1s1*, *igiv2s1_2*, *ighm*, *ighz*), BCR (*cd79b*, *syk*, *lyn*, *blnk*), and other B-lineage genes (e.g., *mpeg1.1*; Figure 4D), but none expressed *cd4* and only ∼14% *cd8*, implying GFP^lo^ thymocytes are T-progenitors just initiating TCR recombination. Conversely, most GFP^hi^ thymocytes expressed *cd8* and/or *cd4* (∼44% DP, ∼41% *cd8^+^*SP, ∼6% *cd4^+^*SP; Figure S3B), *rag1*/*2*, and other T-genes; 30% showed BiP features (Figure 4D, 2^nd^-column). We anticipated scale GFP^lo^ cells would resemble P4. Instead, they exhibited three profiles: 38% were GFP^lo^ thymic BiP cells, 50% T-lineage (like P4), and 12% B-lineage (Figure 4D, 3^rd^-column). Marrow GFP^lo^ cells were similar (45% BiP, 42% T, 10% B; Figure 4D, 4^th^-column).

Comparing both lines, we draw five conclusions: (1) P1 is B-lineage. (2) P2-P4 are largely T-lineage, with P2 (mCh^hi^), P3 (also mCh^hi^), and P4 (mCherry^lo^) all containing BiP lymphocytes. (3) Surprisingly, T-lineage cells are more uniform in dual-transgenic (mCh^lo^ and mCh^hi^ cells in every tissue) than *lck:GFP* (GFP^hi^ cells only in thymus). Cell profiles in *lck*:*GFP* a lso varied more. (4) Like preceding genotypes, B and T lymphocytes are abundant in epidermis. (5) Reproducibly, BiP cells are found in every tissue and genotype.

### Epidermal lymphocyte quantification

Defining cell subtypes and frequencies allowed lymphocyte quantification post-biopsy, via extrapolated identities from gene expression profiles (GEP; Figures 3-4). *cd79a:GFP* scales had ∼40 GFP^hi^ (84% B-lineage, 16% BiP; Figure 3A) and ∼58 GFP^lo^ (50% T, 37% BiP, 8% B) cells/scale (Figure 5A, Table 2). *cd79b:GFP* scales had ∼6 GFP^hi^ (100% B; Figure 3C) and ∼29 GFP^lo^ (50% B, 22% T, 9% BiP) cells/scale. We also tested *IgM1:GFP* (which labels *ighm*^+^ B cells) scales,^14^ finding ∼10 GFP^+^ B cells/scale, less than *cd79* fish, where both Ig-lineages express GFP.

**Figure 5:**
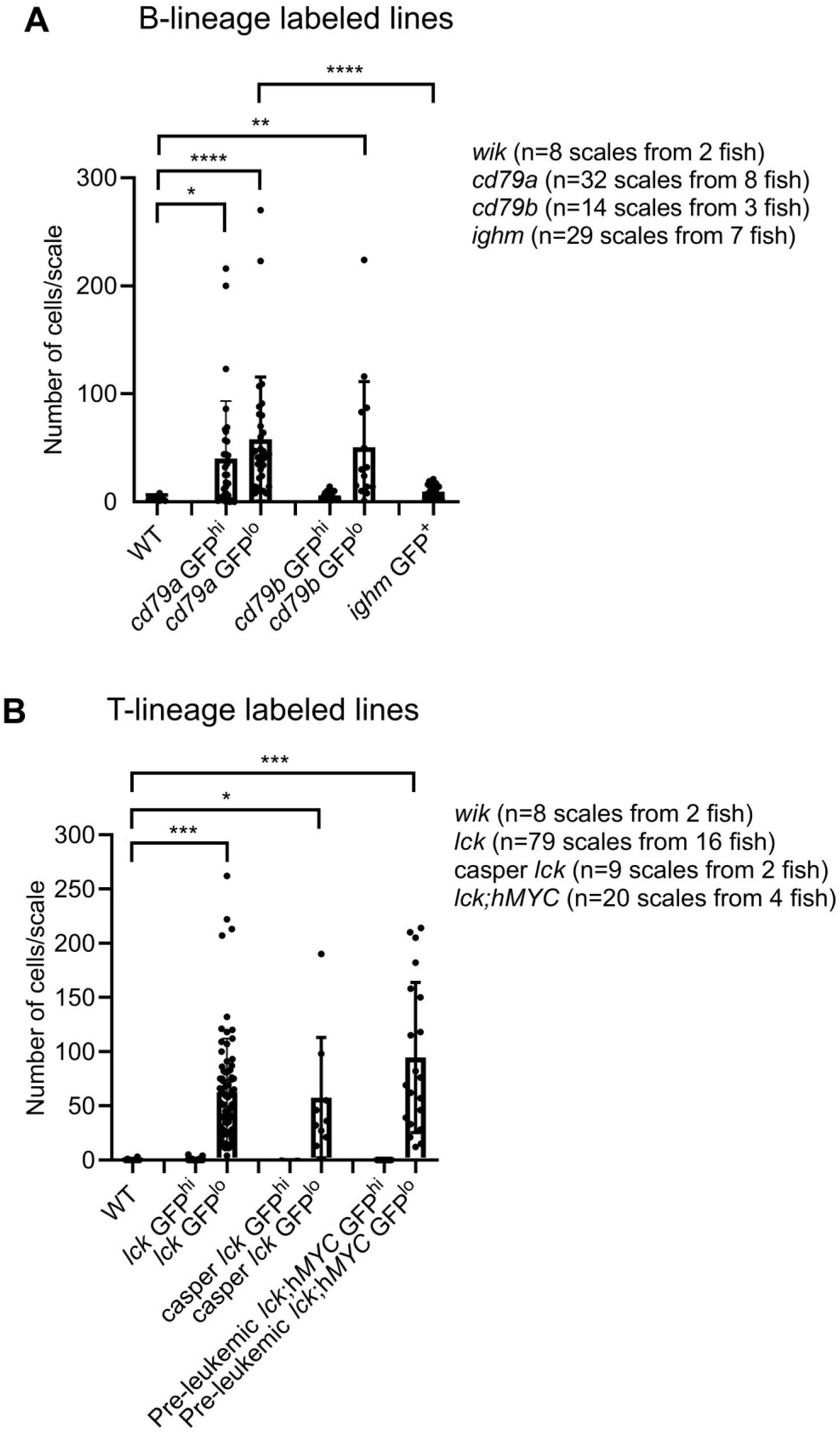
Quantifying scale epidermal lymphocytes. FAC-sorted lymphocytes recovered post-biopsy of **(A)** B- or **(B)** T-lineage labeled lines. WT scales (*WIK* genotype) were auto-fluorescence controls. Bars indicate number of GFP^hi^ or GFP^lo^ cells/scale. Each dot is one scale. Pairwise comparisons to identify significantly different pairs of means were performed using Dunn’s test. *p*-values: *<0.05, **<0.005, ***<0.0005, ****<0.0001.

**Table 2:** Quantification of scale lymphocytes.

| Genotype | *cd79a:GFP* |  | *cd79b:GFP* |  | *IgM1:GFP* | *WIK lck:GFP* |  | *Casper lck:GFP* |  | *lck:GFP + rag2:hMYC* |  |
| --- | --- | --- | --- | --- | --- | --- | --- | --- | --- | --- | --- |
| Fluorescence | GFP<sup>hi</sup> | GFP<sup>lo</sup> | GFP<sup>hi</sup> | GFP<sup>lo</sup> | GFP<sup>+</sup> | GFP<sup>hi</sup> | GFP<sup>lo</sup> | GFP<sup>hi</sup> | GFP<sup>lo</sup> | GFP<sup>hi</sup> | GFP<sup>lo</sup> |
| **B cells/scale** | 34<br>(84%) | 5<br>(8%) | 6<br>(100%) | 15<br>(50%) | 10<br>(100%) |  | 8<br>(12%) |  | 7<br>(12%) |  | 14<br>(15%) |
| **T cells/scale** |  | 29<br>(50%) |  | 6<br>(22%) |  |  | 32<br>(50%) |  | 29<br>(50%) |  | 41<br>(43%) |
| **BiP cells/scale** | 6<br>(16%) | 21<br>(37%) |  | 3<br>(9%) |  |  | 24<br>(38%) |  | 22<br>(38%) |  | 30<br>(32%) |
| **GFP<sup>hi</sup> or GFP<sup>lo</sup> cells/scale** | 40 ± 9 | *58 ± 10 | 6 ± 1 | *29 ± 9 | 10 ± 1 |  | 63 ± 6 |  | 58 ± 19 |  | *95 ± 16 |
| **Lymphocytes/scale** | **98** |  | **35** |  | **10** | **63** |  | **58** |  | **95** |  |

*lck:GFP* scales (Figure 5B, Table 2) had no GFP^hi^ and ∼63 GFP^lo^ cells/scale (50% T, 38% BiP, 12% B; Figure 4D). We also analyzed *lck:GFP* in *casper* (lacking melanophores and iridophores^41^) and *hMYC* backgrounds. *Casper* scales had no GFP^hi^ and ∼58 GFP^lo^ cells/scale, comparable to *WIK lck:GFP*. *hMYC* scales resembled *WIK lck:GFP*, with no GFP^hi^ and ∼95 GFP^lo^ cells/scale (43% T, 32% BiP, 15%B; Figure S3C). A *rag2* promoter drives *hMYC*, and ultimately induces ALL.^26,27,42^ Scales of this genotype had more lymphocytes, but we exclused fish with ALL, and the ∼50% increase (95 vs. 63) was statistically insignificant. Excluding *hMYC* fish, we recovered an average of ∼64 lymphocytes/scale from the *cd79* and *lck* genotypes analyzed (Table 2).

### Lymphocyte scRNAseq

We next performed scRNAseq on GFP^hi^ thymocytes and GFP^lo^ scale/marrow cells from *lck:GFP* fish. After QC, we obtained 1,890 cell GEPs, which clustered into 13 populations (0-12; Figure 6A; analysis details in Supplemental Material). To identify groups, we examined T-/B-/NK-lineage defining genes,^26,43–45^ plus our sc-qPCR data (Supplementary Table 2 genelist). We also examined ILC-genes, but found little expression. UMAP demonstrated the lower cluster group (Populations 3, 5, 6, 8, and 10) was near-exclusive to thymus (Figures S4C-D), suggesting these are immature T cells. Only Population 12 in this lower group (black in Figures 6A, S4A) contained appreciable non-thymocytes, with >1/3 marrow-derived (Figure S4C-D). Relative enrichment scores were to build violin plots, showing Population 3 with the strongest T-lineage profile (dark-green in Figure 6A). Like Population 3, other thymus-derived clusters (Populations 5, 6, 8, 10) had high *rag1/2*, *lck*, *GFP*, and other T-lineage transcripts (Figure S4B). Populations 3, 5, 6, and 8 contained *cd4*^+^/*8*^+^ DP cells, while Population 10 was DN *rag1*/*gata3*^+^. Only Population 8 had little *il7r*. Mammalian DN4 and DP T-lymphoblasts down-regulate *IL7R*, arguing thymic-derived Populations 3, 5, 6, and 10 are likely immature T cells of these stages, with Population 8 being an earlier developmental stage.^46^ A key feature differentiating Populations 5, 6, and 8 was cell-cycle stage: Population 8 was near-entirely S-phase, Population 5 was G_2_/M, and Population 6 contained G_0_/G_1_, S, and G_2_/M cells.

**Figure 6:**
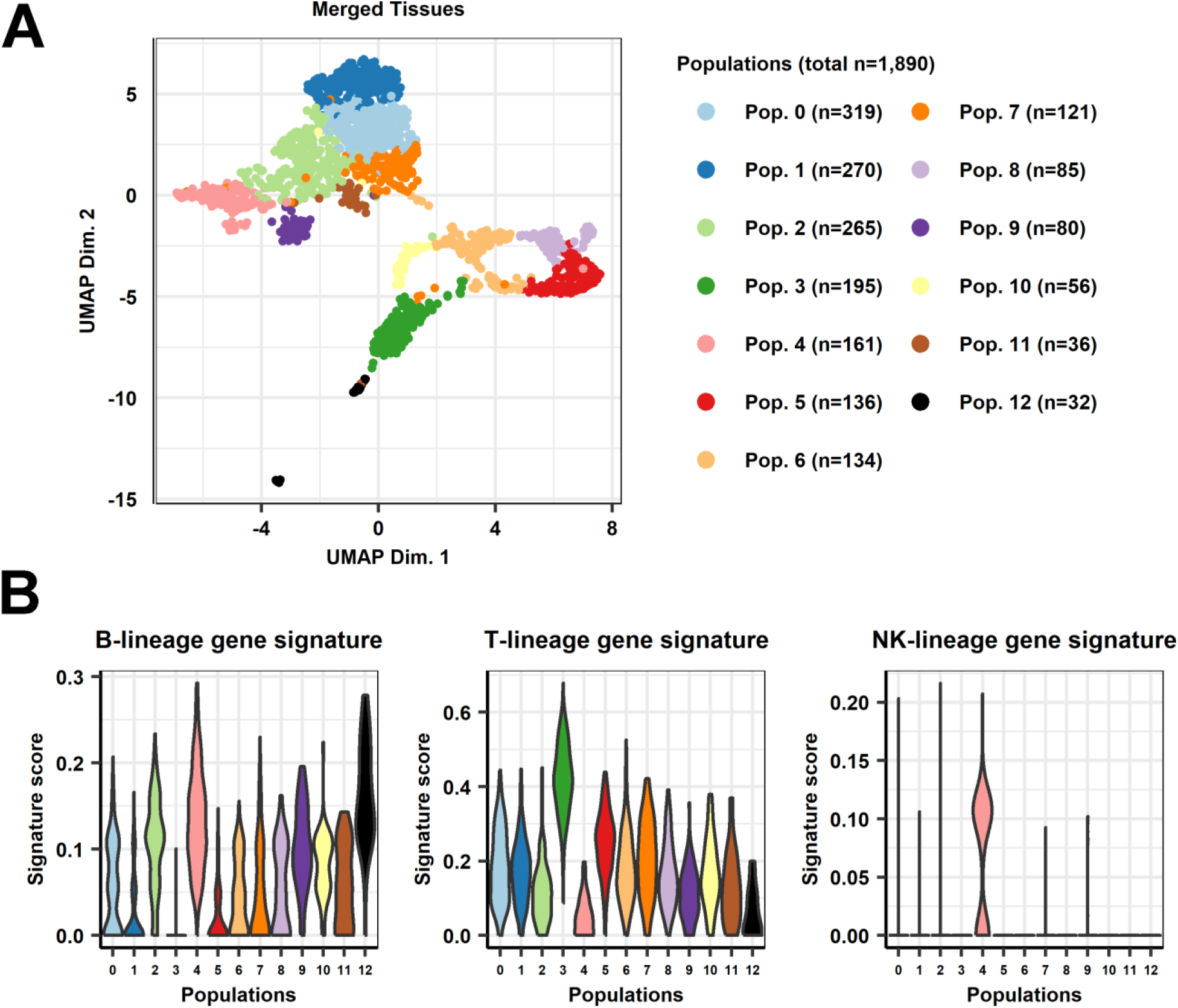
scRNAseq and lineage-specific gene signatures of *lck:GFP* lymphocytes. **(A)** UMAP plot of GFP^+^ cells from thymus, scale, and marrow (merged) from *lck:GFP* fish. Cluster analysis showed 13 populations; cell number for each is listed at right. **(B)** Violin plot distributions of T-, B-, and NK-lineage single-cell gene signature scores for each cluster (estimated using Seurat and UCell). Violin plots are color-coded according to UMAP and legend colors above.

Population 12 (black cluster in Figure 6A) showed a violin plot B-signature, was *cd4/8/rag*^-^, and from only thymus and marrow (Figures S4C-D). Population 12 cells also expressed myeloid markers like *spi1b*, *cepba,* and *fcer1gl* (Supplemental Table 3). Comparison to human genesets by over-representation analysis (details in Supplemental Material) corroborated this, suggesting Population 12 contains primarily myeloid/dendritic cells.

Most upper UMAP clusters (Populations 0, 1, 2, 4, 7, 9; Figure 6A) were seen in all three tissue types, including scales (Figure S4C-D), with only Population 11 absent in scales. Populations 0 and 1 (light- and dark-blue) had violin plot T-lineage signatures and were either *cd4*^+^SP (the majority) or *cd8*^+^SP (the minority; Figure S4B). Population 7 (dark-orange) likewise contained *rag*^-^ mature *cd4*^+^ SP or *cd8*^+^ SP T cells. Populations 2 and 4 (light-green and pink, Figure 6A) were large clusters in the upper UMAP group, most abundant in scales (Figure S4C). Their violin plots indicated B-signatures (Figures 6A, S4D), but many B-specific markers (i.e., *cd79*) were not detected by scRNAseq, except *syk* (both populations) and *lyn* (primarily Population 4; Figure S4B). Much of Population 4 expressed *nkl.2* (a human *GNLY* orthologue; Figure S4B) suggesting it may contain NK cells. Violin plots for Population 4 support this. Over-representation analysis of Population 2 and 4 markers and human genesets indicate both contain NK cells, with Population 2 cells being *id3*^+^.

As noted, Population 11 (brown, Figure 6A) was absent in scales (Figure S4C) and rare overall. Some cells were faintly *cd4*^+^*/8*^+^ DP, but violin plots were inconclusive. Cluster-specific markers *egr2b*, *cebpd*, and *lcp2a,* and high *cd63* levels indicate Population 11 likely includes granulocytes (Supplemental Table 3), which over-representation analysis with human genesets corroborated. Finally, Population 9 cells (purple, Figure 6A) were rare and nearly scale-exclusive (Figure S4C). They were *cd4*^-^*/8*^-^ and *rag1*^-^*/2*^-^, with high *syk* and *il7r* (Figure S4B). Over-representation analysis indicated Population 9 correlated with NK/NKT cells, and this cluster had many differentially-expressed NK-lineage genes (s*la2, cebpb, socs1a*, etc.; Supplemental Table 3). Population 9 resembles Populations 2 and 4, with higher *il7r* being a key difference (Figure S4B).

### Lymphocyte DXM-sensitivity

To functionally test epidermal lymphocytes, we used DXM,^27^ which depletes mammalian lymphocytes.^47,48^ We serially-biopsied fish pre-/post-10d of treatment, analyzing by microscopy and FACS. To exclude potential biopsy-induced inflammatory effects, we alternated flanks for pre-/post-treatment sampling and also serially-biopsied untreated fish.

Following DXM, microscopic fluorescence declined ∼50% in *cd79a* and *cd79b* fish (Figure 7A-B, upper-rows), with ∼85% fewer GFP^hi^/GFP^lo^ scale cells by FACS (Figures 7A-B, lower-rows). *lck:GFP* fish exhibited ∼20% fainter thymic fluorescence and ∼70% fewer scale GFP^lo^ cells (Figure 7C). *lck:mCherry* and double-transgenic (*cd79a:GFP*/*cd79b:GFP* + *lck:mCherry*) fish yielded similar results (Figure 7D-F). Untreated controls had non-significant differences in all parameters (Figure S5A-S5C). We conclude DXM depletes zebrafish epidermal lymphocytes, as in mammals.^47,48^

**Figure 7:**
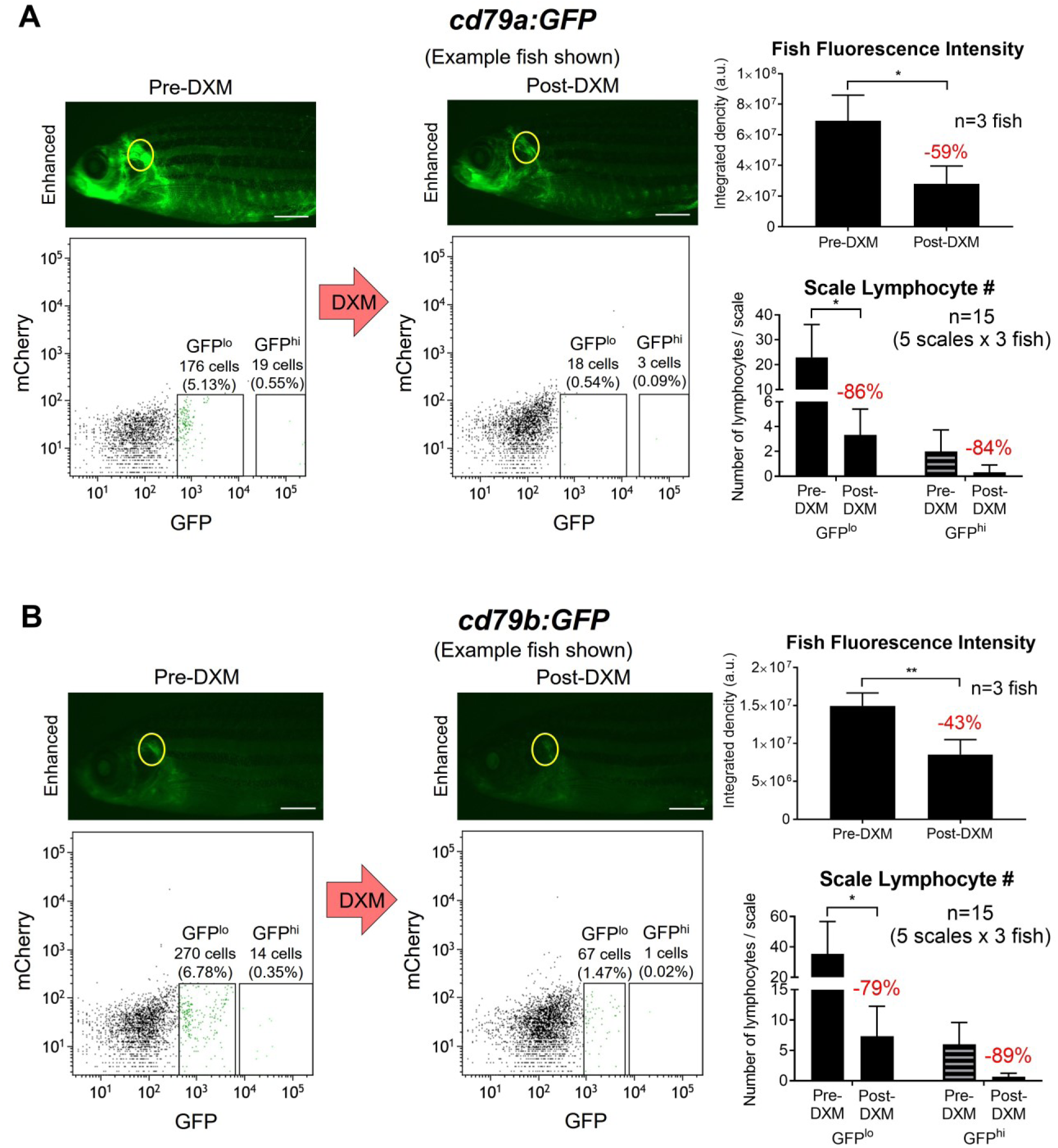
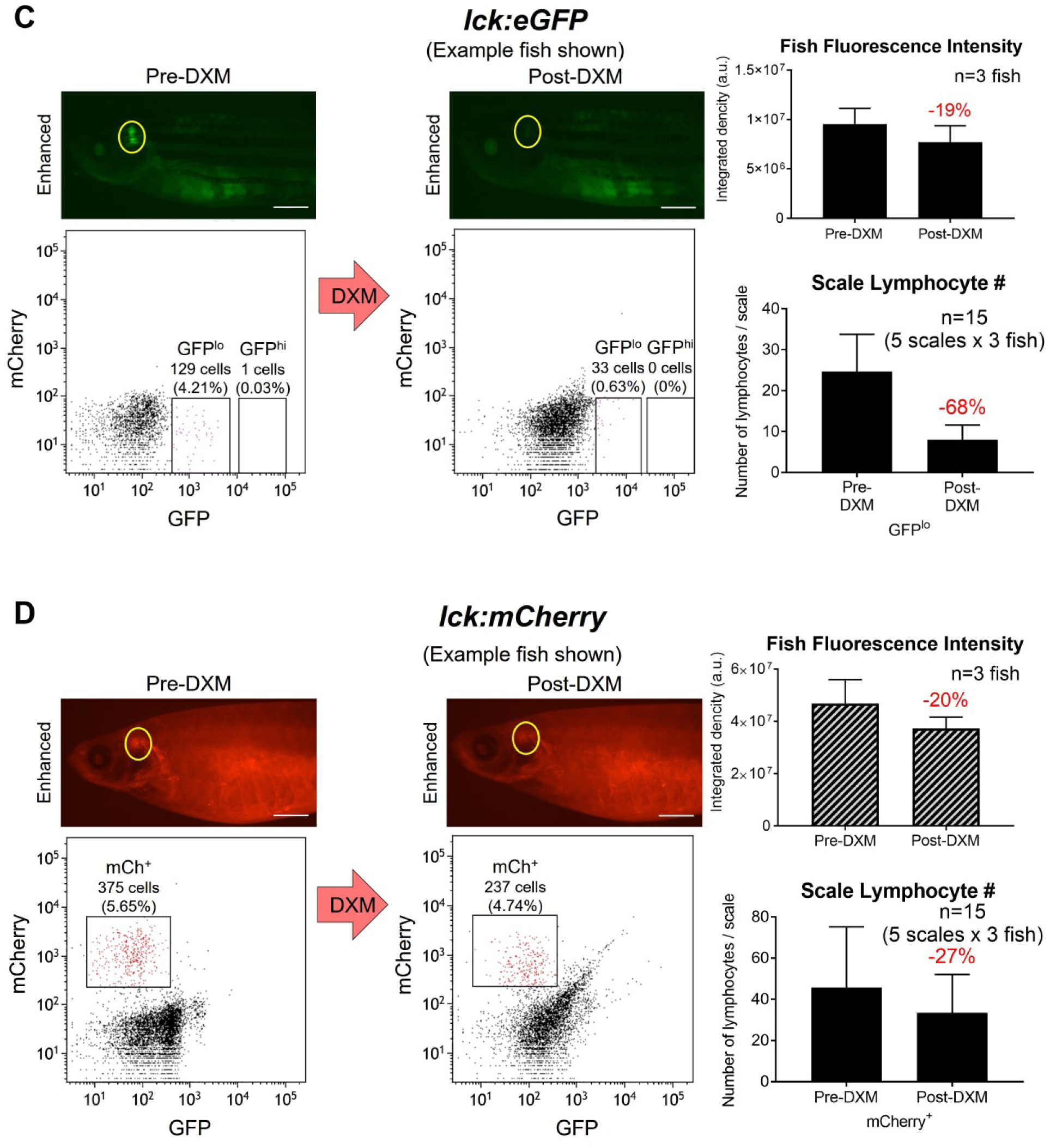
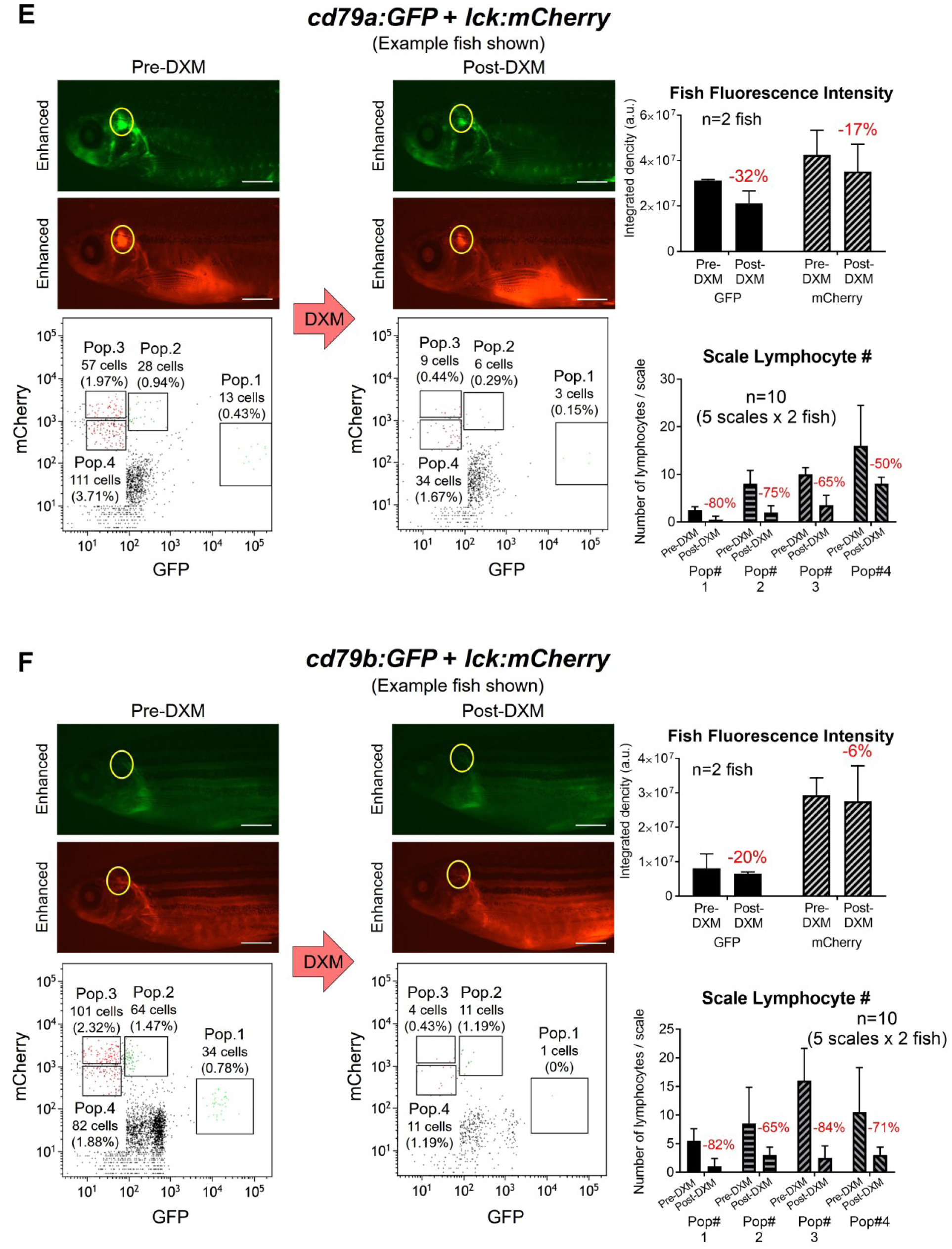
Epidermal lymphocyte DXM responses. (**A**) *cd79a:GFP*, (**B**) *cd79b:GFP*, (**C**) *lck:GFP*, (**D**) *lck:mCherry*, (**E**) *cd79a:GFP* + *lck:mCherry*, and (**F**) *cd79b:GFP* + *lck:mCherry* fish before and after 10 days of continuous DXM immersion. Top panels: pre- and post-DXM fluorescent microscopy images; fluorescence-integrated density histograms at right. Yellow ovals denote thymic region; white bars = 2mm. Bottom panels: flow cytometry plots of biopsies pre- and post-DXM (5 scales/biopsy), with histograms of lymphocytes/5 scales at right. Gates, cells/gate, and % of total cells are shown in flow plots. Three (**A**-**D**) or two (**E**, **F**) fish tested for each genotype. Repeated measures two-way ANOVAs used Sidak’s multiple-comparison test. *p*-values: *<0.05, **<0.005, ***<0.0005, ****<0.0001.

### Lymphocyte phagocytosis

Zebrafish *mpeg1.1*^+^ B cells are phagocytic,^19,21^ and many cells were *mpeg1.1*^+^ (Figures 3-4), so we performed bead-ingestion assays (Figure 8A-C).^19^ *cd79a* GFP^lo^ marrow (81% B; ∼38% *mpeg1.1*^+^; Figure 3A) and *cd79b* GFP^hi^ thymic/scale (93-100% B, >84% *mpeg1.1*^+^; Figure 3C) cells had substantial (22-35%) phagocytosis. In *lck:GFP* fish, GFP^lo^ cells from every tissue and GFP^hi^ thymocytes were highly phagocytic also (26-32% and 59%, respectively). To compare cell types, we categorized groups as B, T, or mixed (B, T, and/or BiP cells) based on prior sc-qPCR. While every category showed phagocytic activity, we saw no significant differences (Figure 8C). Thus, many zebrafish lymphocytes—including those in epidermis—are phagocytic.

**Figure 8:**
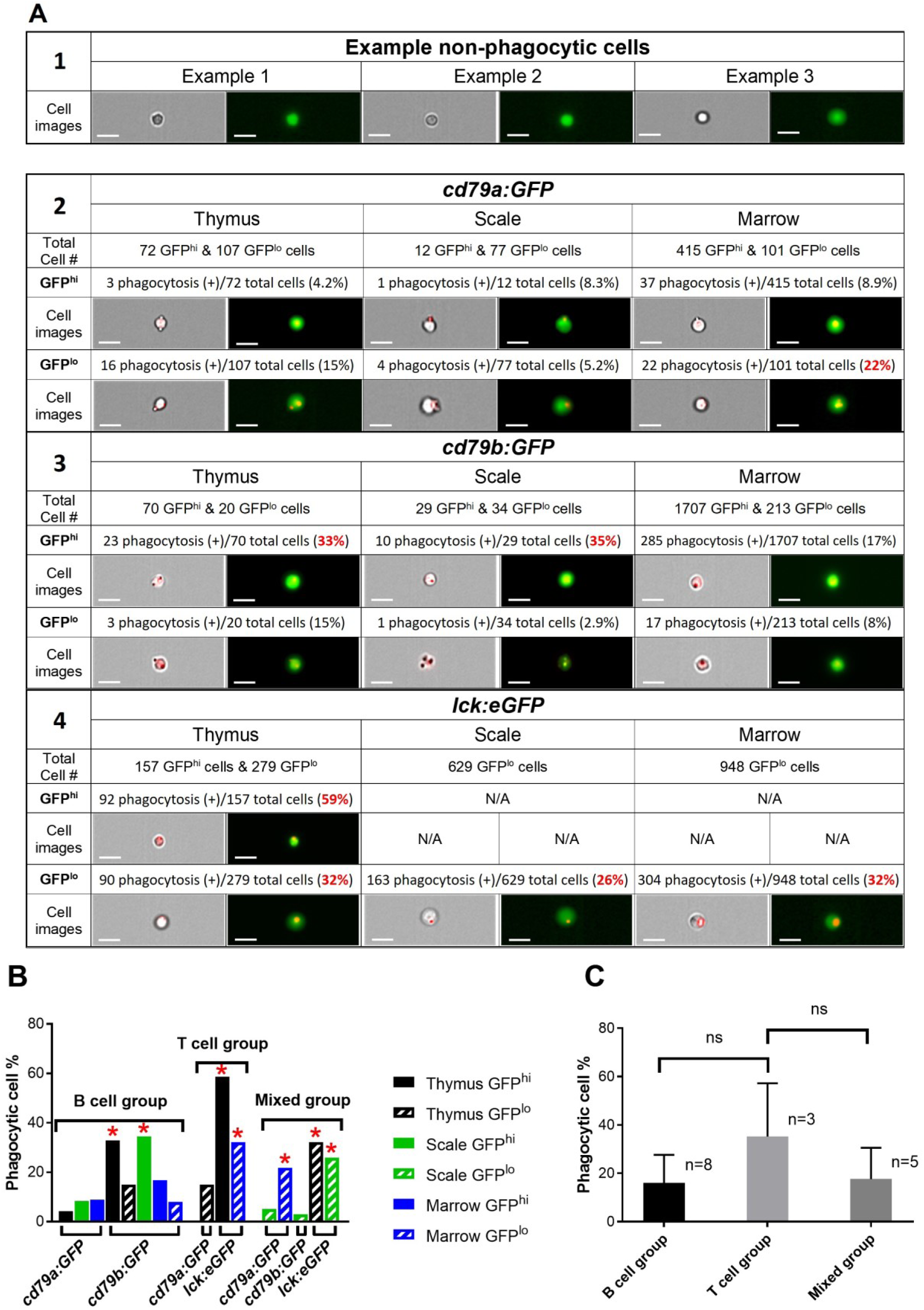
Phagocytosis by zebrafish lymphocytes. **(A)** Results for lymphocytes (GFP^hi^ or GFP^lo^) from each tissue (thymus, scale, or marrow), with total cells tested and percent (+) for red-bead ingestion. Brightfield images show example cells with ≥1 attached/internalized bead. Fluorescent micrographs show merged GFP + red (bead) images. White bars = 10μm. Red font indicates samples with highest phagocytic activity. **(B)** Histograms of phagocytosis results in panel A, grouped by predominant cell type. Asterisks denote samples labeled red in panel A. **(C)** Comparison of each cell type (B, T, or mixed B + T). Groups are based on single-cell expression results in Figures 3-4.

## Discussion

Unlike zebrafish, mammalian epidermal lymphocytes are well-described.^1–5^ Studies in fish have discovered new molecules and cell types,^21,49–51^ and can also inform cutaneous immunity. Here, we provide the first thorough description of zebrafish epidermal lymphocytes. Imaging transgenic fish scales identified epidermal fluorescent cells (Figures 1B, S1B). FACS corroborated this, with GFP^hi^/GFP^lo^ cells in *cd79:GFP* scales and GFP^lo^ cells in *lck:GFP* scales (Figure 2B). qRT-PCR showed GFP^+^ cells in *cd79* lines are B-lineage and GFP^lo^/mCherry^lo^ cells in *lck:fluorophore* fish are T-lineage (Figure 2C).

Single-cell qRT-PCR of lymphocytes from *cd79a* tissues revealed heterogeneity, with most GFP^hi^ cells B-lineage, but GFP^lo^ profiles varying, especially in scales. BiP cells that express T-/B-lineage transcripts equally were detected in all tissues, being most abundant in epidermis (Figure 3A). Their lower frequency in lymphopoietic organs argues they are not lymphoid precursors, and warrants future investigation of this novel population.

Results in *cd79b* fish further clarified cell identities: First, immature and mature thymic B-populations were each detected (Figure 3C). Second, all tissues contained B-lineage GFP^lo^ cells lacking *pax5*, plus other differences from *pax5*^+^ B cells. Our data expand upon a prior *cd79* study,^15^ which noted GFP^-^ *lck^+^* marrow cells (here, GFP^lo^), but did not examine them. Thus, the B-fraction schema^15^ they proposed omitted populations newly described here. Our discovery of T and BiP cells with low *cd79a*/*b* levels further expands both lines’ utility.

We also examined well-characterized *lck*:*GFP* fish^13,16–18,45,52^ Previously, we found GFP^lo^ B-ALL in these^18,26,27^ indicating GFP^lo^ cells were B-lineage, but here GFP^lo^ thymocytes had multiple identities, with individual skin/marrow GFP^lo^ cells having T-, BiP-, and B-GEPs (Figure 4D). Thus, *lck* expression, often considered T-specific, actually varies by tissue, and even within tissues.

We also analyzed double-transgenic fish (Figure 4A-B). P1 (GFP^hi^-only) was uniformly B-lineage and similar across tissues. P2 (GFP^lo^/mCh^hi^) was chiefly T-lineage, but also contained BiP cells in each tissue. P3 (GFP^-^/mCh^hi^) was also primarily T-lineage, but despite no GFP-fluorescence and undetectable *cd79a*, many P3 cells had BiP features, with high *cd79b* and other B-lineage—including *GFP*—transcripts. P4 (GFP^-^/mCh^lo^) resembled P2 and P3, containing both T and BiP cells, but unlike P2-P3, also harbored rare B cells.

Biopsies yielded ∼60 lymphocytes/scale (Figure 5, Table 2). With ∼700 scales and rapid regrowth,^22^ twenty zebrafish scales can provide >10^3^ lymphocytes. Exploiting this, we piloted scRNAseq using *lck*:*GFP* scale cells and lymphopoietic controls. GEP revealed 13 clusters (Figure 6A), with Populations 3, 5, 6 and 8 representing *il7r^-^* DN4/DP T-lymphoblasts, Population 4 exhibiting strong B- and NK-GEP, and Population 2 potentially harboring BiP lymphocytes (Figures 6B, S4B, S4D). Expectedly, scRNAseq of *lck*:*GFP* lymphocytes fails to answer many questions, as this transgene does not label all lymphocytes, even when parsed into GFP^lo^ and GFP^hi^ fractions. scRNAseq of *cd79a*, *cd79b*, and our novel double-transgenic lines will likely resolve these issues.

Functionally, we tested DXM (which depletes mammalian lymphocytes),^47,48^ quantifying epidermal lymphocyte responses. All genotypes showed diminished fluorescence by microscopy and decreased cell numbers by cytometry (Figure 7). Finally, we also demonstrated phagocytosis by multiple lymphocyte populations (Figure 8).

Overall, we describe new methodology to collect fish epidermal lymphocytes, and the first comprehensive description of these cells, advancing zebrafish adaptive immunity modeling. This will facilitate lymphocyte-based investigations, including longitudinal studies where serial-sampling bolsters experimental design.

## Acknowledgments

We thank Aya Ludin and Len Zon (Harvard University) for sharing Tg(*lck:mCherry*) transgenic fish.

## Role of Funding Sources

Studies were supported by grants from the OUHSC Stephenson Cancer Center Pilot Grant Program, the Oklahoma Center for Adult Stem Cell Research, the Presbyterian Health Foundation Team Science and Seed Grant Programs, and the W.J. Jones Family Foundation. Studies by the OUHSC Stephenson Cancer Center Molecular Biology and Cytometry Research Core (confocal microscopy, flow cytometry/FACS, ImageStream™) were supported by NCI Cancer Center Support Grant Award (P30 CA225520).

## Authorship Contributions

GP and JKF conceived and designed the study. GP, MMP, AH, and JJM performed experiments. GP, CAF, JKF analyzed results. GP, CAF, and JKF wrote the manuscript, which all authors approved.

## Declaration of interest

The authors report no conflicts of interest.

## Notes

### Competing Interest Statement

The authors have declared no competing interest.

## References

1. Egawa G, Kabashima K. Skin as a peripheral lymphoid organ: revisiting the concept of skin-associated lymphoid tissues. J Invest Dermatol. Nov 2011;131(11):2178–85. doi:10.1038/jid.2011.198

2. Matejuk A. Skin Immunity. Arch Immunol Ther Exp (Warsz). Feb 2018;66(1):45–54. doi:10.1007/s00005-017-0477-3

3. Ono S, Kabashima K. Proposal of inducible skin-associated lymphoid tissue (iSALT). Exp Dermatol. Aug 2015;24(8):630–1. doi:10.1111/exd.12716

4. Salmi M, Jalkanen S. Lymphocyte homing to the gut: attraction, adhesion, and commitment. Immunol Rev. Aug 2005;206:100–13. doi:10.1111/j.0105-2896.2005.00285.x

5. von Andrian UH, Mempel TR. Homing and cellular traffic in lymph nodes. Nat Rev Immunol. Nov 2003;3(11):867–78. doi:10.1038/nri1222

6. Egbuniwe IU, Karagiannis SN, Nestle FO, Lacy KE. Revisiting the role of B cells in skin immune surveillance. Trends Immunol. Feb 2015;36(2):102–11. doi:10.1016/j.it.2014.12.006

7. Nestle FO, Di Meglio P, Qin JZ, Nickoloff BJ. Skin immune sentinels in health and disease. Nat Rev Immunol. Oct 2009;9(10):679–91. doi:10.1038/nri2622

8. Pasparakis M, Haase I, Nestle FO. Mechanisms regulating skin immunity and inflammation. Nat Rev Immunol. May 2014;14(5):289–301. doi:10.1038/nri3646

9. Streilein JW. Skin-associated lymphoid tissues (SALT): origins and functions. J Invest Dermatol. Jun 1983;80 Suppl:12s–16s. doi:10.1111/1523-1747.ep12536743

10. Sunyer JO. Evolutionary and functional relationships of B cells from fish and mammals: insights into their novel roles in phagocytosis and presentation of particulate antigen. Infect Disord Drug Targets. Jun 2012;12(3):200–12. doi:10.2174/187152612800564419

11. Hansen JD, Landis ED, Phillips RB. Discovery of a unique Ig heavy-chain isotype (IgT) in rainbow trout: Implications for a distinctive B cell developmental pathway in teleost fish. Proc Natl Acad Sci U S A. May 10 2005;102(19):6919–24. doi:10.1073/pnas.0500027102

12. Danilova N, Bussmann J, Jekosch K, Steiner LA. The immunoglobulin heavy-chain locus in zebrafish: identification and expression of a previously unknown isotype, immunoglobulin Z. Nat Immunol. Mar 2005;6(3):295–302. doi:10.1038/ni1166

13. Langenau DM, Ferrando AA, Traver D, et al. In vivo tracking of T cell development, ablation, and engraftment in transgenic zebrafish. Proc Natl Acad Sci U S A. May 11 2004;101(19):7369–74. doi:10.1073/pnas.0402248101

14. Page DM, Wittamer V, Bertrand JY, et al. An evolutionarily conserved program of B-cell development and activation in zebrafish. Blood. Aug 22 2013;122(8):e1–11. doi:10.1182/blood-2012-12-471029

15. Liu X, Li YS, Shinton SA, et al. Zebrafish B Cell Development without a Pre-B Cell Stage, Revealed by CD79 Fluorescence Reporter Transgenes. J Immunol. Sep 1 2017;199(5):1706–1715. doi:10.4049/jimmunol.1700552

16. Moore FE, Garcia EG, Lobbardi R, et al. Single-cell transcriptional analysis of normal, aberrant, and malignant hematopoiesis in zebrafish. J Exp Med. May 30 2016;213(6):979–92. doi:10.1084/jem.20152013

17. Tang Q, Iyer S, Lobbardi R, et al. Dissecting hematopoietic and renal cell heterogeneity in adult zebrafish at single-cell resolution using RNA sequencing. J Exp Med. Oct 2 2017;214(10):2875–2887. doi:10.1084/jem.20170976

18. Burroughs-Garcia J, Hasan A, Park G, Borga C, Frazer JK. Isolating Malignant and Non-Malignant B Cells from lck:eGFP Zebrafish. J Vis Exp. Feb 22 2019;(144)doi:10.3791/59191

19. Ferrero G, Gomez E, Lyer S, et al. The macrophage-expressed gene (mpeg) 1 identifies a subpopulation of B cells in the adult zebrafish. J Leukoc Biol. Mar 2020;107(3):431–443. doi:10.1002/JLB.1A1119-223R

20. Weinstein JA, Jiang N, White RA, 3rd, Fisher DS, Quake SR. High-throughput sequencing of the zebrafish antibody repertoire. Science. May 8 2009;324(5928):807–10. doi:10.1126/science.1170020

21. Zhu LY, Lin AF, Shao T, et al. B cells in teleost fish act as pivotal initiating APCs in priming adaptive immunity: an evolutionary perspective on the origin of the B-1 cell subset and B7 molecules. J Immunol. Mar 15 2014;192(6):2699–714. doi:10.4049/jimmunol.1301312

22. Iwasaki M, Kuroda J, Kawakami K, Wada H. Epidermal regulation of bone morphogenesis through the development and regeneration of osteoblasts in the zebrafish scale. Dev Biol. May 15 2018;437(2):105–119. doi:10.1016/j.ydbio.2018.03.005

23. Bergen DJM, Kague E, Hammond CL. Zebrafish as an Emerging Model for Osteoporosis: A Primary Testing Platform for Screening New Osteo-Active Compounds. Front Endocrinol (Lausanne). 2019;10:6. doi:10.3389/fendo.2019.00006

24. Frantz WT, Ceol CJ. From Tank to Treatment: Modeling Melanoma in Zebrafish. Cells. May 22 2020;9(5)doi:10.3390/cells9051289

25. Pugach EK, Li P, White R, Zon L. Retro-orbital injection in adult zebrafish. J Vis Exp. Dec 7 2009;(34)doi:10.3791/1645

26. Borga C, Park G, Foster C, et al. Simultaneous B and T cell acute lymphoblastic leukemias in zebrafish driven by transgenic MYC: implications for oncogenesis and lymphopoiesis. Leukemia. Feb 2019;33(2):333–347. doi:10.1038/s41375-018-0226-6

27. Park G, Burroughs-Garcia J, Foster CA, Hasan A, Borga C, Frazer JK. Zebrafish B cell acute lymphoblastic leukemia: new findings in an old model. Oncotarget. Apr 14 2020;11(15):1292–1305. doi:10.18632/oncotarget.27555

28. Meyer LH, Eckhoff SM, Queudeville M, et al. Early relapse in ALL is identified by time to leukemia in NOD/SCID mice and is characterized by a gene signature involving survival pathways. Cancer Cell. Feb 15 2011;19(2):206–17. doi:10.1016/j.ccr.2010.11.014

29. Brown W, Deiters A. Light-activation of Cre recombinase in zebrafish embryos through genetic code expansion. Methods Enzymol. 2019;624:265–281. doi:10.1016/bs.mie.2019.04.004

30. Traver D, Paw BH, Poss KD, Penberthy WT, Lin S, Zon LI. Transplantation and in vivo imaging of multilineage engraftment in zebrafish bloodless mutants. Nat Immunol. Dec 2003;4(12):1238–46. doi:10.1038/ni1007

31. Bauer TR, Jr., McDermid HE, Budarf ML, Van Keuren ML, Blomberg BB. Physical location of the human immunoglobulin lambda-like genes, 14.1, 16.1, and 16.2. Immunogenetics. 1993;38(6):387–99. doi:10.1007/BF00184519

32. Palmer C, Diehn M, Alizadeh AA, Brown PO. Cell-type specific gene expression profiles of leukocytes in human peripheral blood. BMC Genomics. May 16 2006;7:115. doi:10.1186/1471-2164-7-115

33. Clark MR, Mandal M, Ochiai K, Singh H. Orchestrating B cell lymphopoiesis through interplay of IL-7 receptor and pre-B cell receptor signalling. Nat Rev Immunol. Feb 2014;14(2):69–80. doi:10.1038/nri3570

34. Asnafi V, Beldjord K, Garand R, et al. IgH DJ rearrangements within T-ALL correlate with cCD79a expression, an immature/TCRgammadelta phenotype and absence of IL7Ralpha/CD127 expression. Leukemia. Dec 2004;18(12):1997–2001. doi:10.1038/sj.leu.2403531

35. Garg N, Kotru M, Kumar D, Pathak R, Sikka M. Correlation of expression of aberrant immunophenotypic markers in T-ALL with its morphology: A pilot study. J Lab Physicians. Oct-Dec 2018;10(4):410–413. doi:10.4103/JLP.JLP_35_18

36. Mangogna A, Cox MC, Ruco L, Lopez G, Belmonte B, Di Napoli A. Rituximab Plus Chemotherapy Provides No Clinical Benefit in a Peripheral T-Cell Lymphoma Not Otherwise Specified with Aberrant Expression of CD20 and CD79a: A Case Report and Review of the Literature. Diagnostics (Basel). May 26 2020;10(6)doi:10.3390/diagnostics10060341

37. Borga C, Foster CA, Iyer S, Garcia SP, Langenau DM, Frazer JK. Molecularly distinct models of zebrafish Myc-induced B cell leukemia. Leukemia. Feb 2019;33(2):559–562. doi:10.1038/s41375-018-0328-1

38. Garcia EG, Iyer S, Garcia SP, et al. Cell of origin dictates aggression and stem cell number in acute lymphoblastic leukemia. Leukemia. Aug 2018;32(8):1860–1865. doi:10.1038/s41375-018-0130-0

39. Brennan PJ, Brigl M, Brenner MB. Invariant natural killer T cells: an innate activation scheme linked to diverse effector functions. Nat Rev Immunol. Feb 2013;13(2):101–17. doi:10.1038/nri3369

40. Tindemans I, Serafini N, Di Santo JP, Hendriks RW. GATA-3 function in innate and adaptive immunity. Immunity. Aug 21 2014;41(2):191–206. doi:10.1016/j.immuni.2014.06.006

41. White RM, Sessa A, Burke C, et al. Transparent adult zebrafish as a tool for in vivo transplantation analysis. Cell Stem Cell. Feb 7 2008;2(2):183–9. doi:10.1016/j.stem.2007.11.002

42. Gutierrez A, Grebliunaite R, Feng H, et al. Pten mediates Myc oncogene dependence in a conditional zebrafish model of T cell acute lymphoblastic leukemia. J Exp Med. Aug 1 2011;208(8):1595–603. doi:10.1084/jem.20101691

43. Carmona SJ, Teichmann SA, Ferreira L, et al. Single-cell transcriptome analysis of fish immune cells provides insight into the evolution of vertebrate immune cell types. Genome Res. Mar 2017;27(3):451–461. doi:10.1101/gr.207704.116

44. Garcia-Valtanen P, Martinez-Lopez A, Lopez-Munoz A, et al. Zebra Fish Lacking Adaptive Immunity Acquire an Antiviral Alert State Characterized by Upregulated Gene Expression of Apoptosis, Multigene Families, and Interferon-Related Genes. Front Immunol. 2017;8:121. doi:10.3389/fimmu.2017.00121

45. Hernandez PP, Strzelecka PM, Athanasiadis EI, et al. Single-cell transcriptional analysis reveals ILC-like cells in zebrafish. Sci Immunol. Nov 16 2018;3(29)doi:10.1126/sciimmunol.aau5265

46. Hong C, Luckey MA, Park JH. Intrathymic IL-7: the where, when, and why of IL-7 signaling during T cell development. Semin Immunol. Jun 2012;24(3):151–8. doi:10.1016/j.smim.2012.02.002

47. Giles AJ, Hutchinson MND, Sonnemann HM, et al. Dexamethasone-induced immunosuppression: mechanisms and implications for immunotherapy. J Immunother Cancer. Jun 11 2018;6(1):51. doi:10.1186/s40425-018-0371-5

48. Aston WJ, Hope DE, Cook AM, et al. Dexamethasone differentially depletes tumour and peripheral blood lymphocytes and can impact the efficacy of chemotherapy/checkpoint blockade combination treatment. Oncoimmunology. 2019;8(11):e1641390. doi:10.1080/2162402X.2019.1641390

49. Zhu LY, Nie L, Zhu G, Xiang LX, Shao JZ. Advances in research of fish immune-relevant genes: a comparative overview of innate and adaptive immunity in teleosts. Dev Comp Immunol. Jan-Feb 2013;39(1-2):39–62. doi:10.1016/j.dci.2012.04.001

50. Zou J, Redmond AK, Qi Z, Dooley H, Secombes CJ. The CXC chemokine receptors of fish: Insights into CXCR evolution in the vertebrates. Gen Comp Endocrinol. May 1 2015;215:117–31. doi:10.1016/j.ygcen.2015.01.004

51. Li J, Barreda DR, Zhang YA, et al. B lymphocytes from early vertebrates have potent phagocytic and microbicidal abilities. Nat Immunol. Oct 2006;7(10):1116–24. doi:10.1038/ni1389

52. Kernen L, Rieder J, Duus A, Holbech H, Segner H, Bailey C. Thymus development in the zebrafish (Danio rerio) from an ecoimmunology perspective. J Exp Zool A Ecol Integr Physiol. Dec 2020;333(10):805–819. doi:10.1002/jez.2435

