## Supplemental Figure for "Diverse Epithelial Lymphocytes in Zebrafish Revealed Using a Novel Scale Biopsy Method"

### Supplementary Materials

#### Supplementary Methods

##### Single-cell qRT-PCR analysis

Twenty-cycle pre-amplifications were performed using gene-specific primers (Supplementary Table 1). Unincorporated primers were digested with exonuclease I (NEB M0293L) and samples diluted 5-fold in DNA suspension buffer (TEKnova PN T0221). Single-cell pre-amplified cDNAs were quantified by qRT-PCR using a Fluidigm BioMark HD instrument and 48.48 Dynamic Array Chips for Gene Expression following manufacturer protocol. A Limit of Detection (LoD) of 27 was applied to data.

##### Scale biopsy

50 mL of 1x Roswell Park Memorial Institute Medium (RPMI) containing 1% fetal bovine serum and 1% penicillin-streptomycin (sorting media) was aliquoted into 1.5 mL Eppendorf tubes. 0.02% tricaine (MS-222) in fish system water was used to anesthetize fish. Sedated fish were placed in Petri dishes and screened for fluorescence using a Nikon AZ100 fluorescent microscope and Nikon DS-Qi1MC camera. Scales in the upper $2^{nd}$ or $3^{rd}$ stripes were flipped forward (i.e., caudally) using a 31-gauge syringe and removed with forceps. Scales were collected on forceps tips until 10-20 scales were obtained, and then placed in 1.5 mL tubes with 500 µl of cold sorting media. Scale tissue was manually dissociated using a pestle micro-tube homogenizer to displace epidermal cells from scales, and then filtered through 35 µm nylon mesh filters to make single-cell suspensions. Single-cell suspensions were FAC-sorted using a BD-FACSJazz Instrument (Becton Dickinson, San Jose, CA, USA). Flow cytometry was performed using a CytoFLEX™ instrument and Kaluza Analysis software (Beckman Coulter, Brea, CA, USA).

**Single-cell RNAseq and analysis**

After creating single-cell emulsions, uniquely-identifiable 1st-strand template single-cell cDNA libraries were generated from each cell by emulsion PCR. 2nd-strand cDNA was generated and ligated to compatible Illumina adapters. Libraries were loaded in single NovaSeq6000 lanes, using read lengths of 28 bp for the first read, 120 bp for the second read, and 8 base index reads. After conversion to fastq files, reads were processed and aggregated using the 10x Genomics *Cell Ranger*[1] (v.6.0.0) pipeline (no normalization, default settings) and processed using the *Seurat* R package (v.4.3.0). We obtained transcriptomes for 6,359 cells after Cell Ranger processing. *SoupX*[2] (v.1.6.2) was used to model and remove ambient RNA contamination, and *scDblFinder*[3] (v.1.12.0) used to detect potential multiplets (default settings, per individual tissue type). Additional QC filtering to remove potential dead/dying cells and cells with abnormal read/gene counts and high mitochondrial transcripts yielded the 1,890 cells shown in analyses. Using *fastMNN*,[4] cells were normalized and integrated, and clustered using *Seurat* (Leiden algorithm). Clustering resolution was optimized using the *clustree*[5] package (v.0.5.0). Cluster boundaries were manually examined and fine-tuned to optimize interpretation. Collective diagnostic gene signatures comprised of published gene lists and our sc-qRT-PCR results were explored using *Seurat*[6] and *UCell*[7] *(v.2.2.0)*. Cell types were investigated and inferred using a combination of the previously-described gene signatures, *cd4/cd8/rag1/rag2* transcript abundance, tissue of origin, and unbiased cell-type recognition using putative human orthologs of differentially-expressed cluster markers as input for over-representation analysis using the *clusterprofiler*[8] package (v.4.9.0.002; using the C8 cell type signatures from MSigDB,[9] and genesets from the SaVanT[10] and

CellMarker2.0[11] databases). Preferential markers were determined for each population using the *FindAllMarkers* function in *Seurat*, to aid in cell-type assignment (p.adj ≤ 0.05, min.pct = 0.25).

**Supplementary Table 1**

| | Outer Forward Primer Sequence | Outer Reverse Primer Sequence | Inner Forward Primer Sequence | Inner Reverse Primer Sequence | NCBI Ref. Seq. | Source | Additional information |
|---|---|---|---|---|---|---|---|
| bcl11a a | GGTTGAGAGAGCTTGCTGGT | AGACTTGGTCTTCATGGGCG | CTAATCGGCCCAGCCCTATG | GGACTGTTGGTTGAGAGGGG | NM_001040391.1 | | |
| bcl6a | CATCACAAACACTGCTCGCC | CCAACGTCTCCCGCTTCATA | CGAGTGACGGTAACAACCCA | CCAGCTCCCTGGAGGAGTAT | XM_02147681 5.1 | | |
| pax5 | CAGCACAACACTCCCAGGAT | AATAGTCCCGATGACGCTGC | AAGGCAGTTACTCCACACCC | ACCGTACTCCTGCTGAAACAC | XM_005170923.4 | | |
| cd79a | CCATGGCACCTTCCTTCAAATA | TGGTAAGCTGAGTTGCAGTCAT | CAGCGAGGGTGTGAAAAACA | CCCTTTCTGTCTTCCTGTCCA | NM_001326470.1 | | |
| cd79b | TTGGCGTTAAGACAGGTCGG | GTCGAACCAGAGGAACGACA | TACTGTGTGCCGTCGAATCC | GTGCCACTGTCCTCAGTCTC | XM_021480215.1 | | |
| cxcr4a | CGTGCTTCCCGGGCTCGTGA | ACAGCAGTGAAAGTACGCGA | CACCGTGGTTCTGATCGTCT | GGGGAATCACCTCCAGCATC | NM_131882.3 | | |
| cxcr4b | TTCCGCTTCCAGCACATCAT | GCAGTGGAAATATGCCAGCG | CCGTCATCCTCATCCTCTGC | CAGCGCCTCCGTAAAGAAGA | NM_131834.1 | | |
| IgM (FW1+RV1) | AGCTTCTCTAGCTCCACCAG | ATTTTGGTGAAATGGAATTG | ACGCAGAAACATGGAAGCAG | TTCAGCATGTCTTCAGGGGTG | 378762 | Gene ID | Ensembl version: ENSDARG00000096355.4 |
| IgZ (FW1+RV1) | AAAGCAACGATACCAAAGTG | AACAGCTTGCAAGACAATTC | TGGTCTTCACACAGCACTCA | GGTGGCTTCAGCATCAGTCT | 446139 | Gene ID | Ensembl version: ENSDARG00000096280.3 |
| id3 | AAGCTGCTATGAAGCGGTGT | AATCTCCACTTGGCTCACGG | CTGCAAGAGTCCTTCCGAGG | TGGCTCACGGACTTGTTCTG | NM_152967.1 | Gene ID | Ensembl version: ENSDARG00000054823.5 |
| btk | GGCGGTTTCTTGGTGAGAGA | CAAAACGCCAAATGCCCAGA | ATGTCAGCGAGGGCATGAAA | GACTTCAGGAGGTGACCAGC | XM_021481245.1 | | |

| | | | | | | | |
|---|---|---|---|---|---|---|---|
| Syk | TGAACGACACCTACGCCATC | CCGTTGACCCGTGAGGTATT | AGCTGGAGAAACTCATCGCC | GCAAACGCAGCTCTGAATCC | NM_212843.2 | | |
| Lyn | ACATCCACCGAGACCTGAGA | TTCTGGTGCCGTCCATTTGA | GGTTTCAGAGATGCTGCTGT | TGGCCGTGTATTGGTCATCC | NM_001004543.1 | | |
| blnk | CAACCCAGCAAACAACCCAG | GACTCCTCTGTTCACCACCG | CGAGGACGACTACATCGAGC | GTCAGTGTTCTCTCCGTGGG | NM_212838.1 | | |
| dtx1 | AAGTGGTGGACTGTTGGGTG | AGAGGACTCGGGAAAGGGAA | GTTTCAGGGGTCAGTGGAGG | CCCAAGAGTTTGTGCCATGC | NM_001318952.1 | Gene ID | Ensembl version: ENSDARG00000102042.2 |
| spi1b | CTCACAACGTCCAGCCATCT | TTGCGATTGCCCTTCTGGAT | ATCCCAGCAGTCGTAGTCCT | TCTCGATCCACCCACCAGAT | NM_198062.2 | | |
| cybb | GGTGGCCTACATGATTGCCT | TGCGTGAACCAGAAGACCTC | ACAGCCGTCCACATCATAGC | ATCACGACACCCGTCAATCC | NM_200414.1 | | |
| igic1s1 | GAGCAGCAGTGGATGGAGAG | AGTGTGTGTGCTGAATGCAG | CAGGAGACACTGTGAGGAG | TAGCAGGAGTGTGTGTGTGC | XM_017353889.1 | | |
| igiv2s1_2 | AGCAGCCTGACTCTCTCTGA | ACAGATGTGGTGTGTGTGCT | GAGCAGCAGTGGATGGAGAG | CACAGTGTATTCTGCAGTGT | ZDB-GENE-040112-1 | ZFIN ID | |
| foxo1b | GACCCGGATTTTGAGCCTCT | GTTGTCCTGGCAGTGGAAGT | AAACCGGAGCAAGGAATCGT | GGCTTCTCGTCAGGGTAGTC | NM_001082857.1 | Gene ID | Ensembl version: ENSDARG00000061549.4 |
| lyl1 (LOC100333113) | TCTGAGCTGCGCAAACTGAT | CTCTCCTCGCTGTCAGTGTC | AAGCTGAGCAAGAACGAGAT | GATCCAGAGGATGCGGTCAG | XM_009302066.3 | Gene ID | ZFIN ID: ZDB-GENE-090807-1 |
| si:dkey-24p1.1 (cd22) | CCCATCCACAGCCAGTCTAC | ATTCAGTGCTGTTCCTGCGA | AGAAACTCAGCCAGTGACCA | CTGCGACACTCTGGGATGTT | NM_001098245.1 | Gene ID | Ensembl version: ENSDARG00000039096.8 |
| pik3c3 | GTCTGGAGGGAAAGCGAGAG | TCCACACCCGGCCACACCTT | TGTCAGGTGTTTGCTGAGGG | CAGGTCCATACACGTCCCAC | NM_001328533.1 | | |

| | | | | | | | |
|---|---|---|---|---|---|---|---|
| rasa4 | GATCCGTG CGGTAGAG AAGG | AGATGAGC TGAGCCTC CAGA | TGGAGGA GAAGTGT TTCGGC | TCTGCTTT GTAGATG CCGGG | NM_0 01099 451.1 | | |
| lck-5 | AGATTGCT GACTTCGG CCTG | GTAGCCCC TCTCGAGG TTTG | GGCACCA GAGGCCA TAAACT | CTCTGGG TTTGTCAT TCCTGG | NM_0 01001 596.1 | | |
| cd2 (si:ch2 11- 132g1. 1) | GGTGGAG GAGCTCTT TTGCT | AGACGGGT TACAGTGC ATGG | CGCAAGA AGCACCG ACTAGA | CTTCTTGT GGGATAG GGGGC | XM_00 51683 09.3 | | |
| cd4-3 (cd4- 1) | TGTGTTGC CATCGGGA GTGG | GGCAAACA TTGTGCAG AGAACT | TCTTGCTT GTTGCATT CGCC | TCCCTTTG GCTGTTTG TTATTGT | NM_0 01135 096.1 | | |
| cd8a-3 | TGCAAAAA GGACAGAC AGCG | AAAGTCCA CAACCTCC GACC | ACTCTTCT TCGGAGA GGTGAC | ACAGGCT TCAGTGTT GTTTGAA | NM_0 01040 049.1 | Gen e ID | Ensembl version: ENSDARG000 00044797.5 |
| Notch 1a | AACCTGCC TCGACCAG ATTG | CGTCTGTG CACTTAGC TCCA | CATCTCAG CCGTGTCT CAAC | AACCCTTT AGGGCAT TCGCA | NM_1 31441. 1 | | |
| ets1 | CCTATCAG ACGCTTCA CCCC | AGGCTGTT GAAGGAC GACTG | AAGACCT GCTGTCG CTCAAA | GTCCTGA CCACCCA GCTTAC | NM_0 01017 558.1 | | |
| il7r | GCAAGTAC ACTTAAGA GACGGCA | CCATCTCC ACCAAGTC TGTC | TGGTCAA AATTCCA GCTCCTG A | TGTTCACC ACGGATC TCCAC | NM_0 01113 507.2 | | |
| itk | ATGGAGCC AAACCTCT TCCG | ACCATGAA ACCCCCGT CCTT | CTGGTGG ACCGTGA AAGACA | AAACCCCC GTCCTTGT TCTC | NM_1 31104. 1 | Gen e ID | Ensembl version: ENSDARG000 00017565.8 |
| Lat | AACTGCTG TCTATGGT GGGC | CTGAAGGA GGACAGG AGGGA | CGGCATC CCCATTCG ACATA | CTCAGCG AAGGGTG TATTGG | NM_0 01143 684.1 | | |
| Skap1 | AAAGACCA AGCACTGC CTCA | CCTCTCTG GAAGGCCA ACTC | GCAGTGA GAGAACT GGGTCC | CGGTCCCT CAGCTTCA CAAT | NM_0 01080 011.1 | | |
| tcf7 | GTCAGCGA GACCGAGA TCAG | AAACCCGA AATCTCCT GCGT | CCAACGG ACCCGTCT CTCCA | CCTGGTTT CTGTCCTC CGTC | NM_0 01012 389.2 | | |

| | | | | | | | |
|---|---|---|---|---|---|---|---|
| Zap-70 | TGTGCGCATGATTGGACTCT | CCAGGAGCACATTACGAGCA | CCCGCTCAACAAGTTCCTCT | TCCCCATTGACACCTGATGC | NM_001020589.1 | | |
| Gata3 | AAACGACCACGACGACACTG | GAAAAGGGAGGGAACGAGGTC | ACACAATATCAACCGACCGCT | CCTCCATGCTGTCATGGGAC | NM_131211.1 | Gene ID | Ensembl version: ENSDARG00000015752.9 |
| Mpeg 1.1 | AGACCCACCAAGTGAAAGAGG | CAATGTGGCTCCAGCATCAAC | CGGGTTCAAGTCCGTAACCA | TGGCGTCAGCGATTTCTTCT | NM_212737.1 | Gene ID | Ensembl version: ENSDARG00000055290.5 |
| L-Plastin | CATCCTGGAGGATCTGGGCG | ATCTTACGCGCCATCGAGATT | GCAGTGGGTGAACGAAACAC | CAGCAGGTCGTAGCGGATAG | NM_131320.3 | Gene ID | Ensembl version: ENSDARG00000023188.10 |
| BA1 | TAACCCCAAAGTGGCAGCTC | AGCATTGAAACCAGCTTGGC | GGGAGGTCTTGAGAGAGCCA | GCAATCAGCGAGAAGCCTGA | NM_131020.3 | Gene ID | Ensembl version: ENSDARG00000089087.7 |
| rag1 | CCAGGTGAAGACATTTGCCG | ATTACGCAGAGTGTGCAGGG | AGCAATGATGCAAGGCAGAG | TGTGCAGGGGCTGGAATATC | NM_131389.1 | | |
| rag2 (FW1+RV1) | CGCACTGAATGCTGGGTACTG | AAGACCTAACTACACACTTCGCTT | AATGTGCGTCTCAACGGGAA | TTCTTTCGCTGTTTCCCCGA | NM_131385.3 | Gene ID | Ensembl version: ENSDARG00000052121.6 |
| myca | GTGGTCAGCCAGAGCTTCAT | GCTGGAGCTGTTAGGTGGAG | AAAGGAAGGAACTGATGTCT | ATGGGAAGACCACAGAGGGA | NM_131412.1 | | |
| mycb (FW1+RV1) | GCCTTCTTCTTCACAGGCACT | TAACCATCAGCACGGACAGC | TGCCGCTGAATTCAAGTATGG | TTTGACAACGAGGACGAGGA | NM_200172.1 | Gene ID | Ensembl version: ENSDARG00000007241.8 |
| hmyc | CGTCCTCGGATTCTCTGCTC | GCTGGTGCATTTTCGGTTGT | CAGCGACTCTGAGGAGGAAC | GCTGCGTAGTTGTGCTGATG | NM_002467.5 | | |
| GFP | GACTTCAAGGAGGACGGCAA | TCTCGTTGGGGTCTTTGCTC | ATGGCCGACAAGCAGAAGAA | CTCAGGTAGTGGTTGTCGGG | AAB02572.1 | GenBank | |
| EF1a | AGCGTGGTATCACCATTGACA | TTCACTCCCAGGGTGAAAGC | GAGACCAGCAAATACTACGTC | GGAGATACCAGCCTCAAACTC | NM_131263.1 | | |

| | | | | | |
|---|---|---|---|---|---|
| rpl13a | AGGCTGAA GGTGTTTG ATG | TTTCAGAC GCACAATC TTGA | AGGTGTT TGATGGC ATCCCT | ATCTTGA GAGCAGC TGGGAC | NM_2 12784. 1 |

**Supplementary Table 2**

| gene.name | group | common.name | ensembl ID |
|---|---|---|---|
| bcl11aa | B cell | bcl11aa | ENSDARG00000061352 |
| cd79a | B cell | cd79a | ENSDARG00000037473 |
| cd79b | B cell | cd79b | ENSDARG00000104691 |
| ighm | B cell | ighm | OTTDARG00000004496 |
| btk | B cell | btk | ENSDARG00000004433 |
| syk | B cell | syk | ENSDARG00000008186 |
| lyn | B cell | lyn | ENSDARG00000067916 |
| blnk | B cell | blnk | ENSDARG00000042722 |
| CABZ01004876.1 | B cell | dtx1 | ENSDARG00000102042 |
| spi1b | B cell | spi1b | ENSDARG00000000767 |
| cybb | B cell | cybb | ENSDARG00000056615 |
| igic1s1 | B cell | igic1s1 | ENSDARG00000093272 |
| foxo1b | B cell | foxo1b | ENSDARG00000061549 |
| CABZ01066694.1 | B cell | lyl1 | ENSDARG00000110178 |
| mpeg1.1 | B cell | mpeg1.1 | ENSDARG00000055290 |
| cd4-1 | T cell | cd4-1 | ENSDARG00000070668 |
| cd4-2.2 | T cell | cd4-2.2 | ENSDARG00000073950 |
| cd8a | T cell | cd8a | ENSDARG00000044797 |
| cd8b | T cell | cd8b | ENSDARG00000058682 |
| pik3c3 | T cell | pik3c3 | ENSDARG00000054829 |
| rasa4 | T cell | rasa4 | ENSDARG00000029372 |
| lck | T cell | lck | ENSDARG00000102525 |
| il7r | T cell | il7r | ENSDARG00000078970 |
| itk | T cell | itk | ENSDARG00000017565 |
| BX784026.1 | T cell | lat | ENSDARG00000100334 |
| skap1 | T cell | skap1 | ENSDARG00000001498 |
| tcf7 | T cell | tcf7 | ENSDARG00000038672 |
| zap70 | T cell | zap70 | ENSDARG00000015752 |
| gata3 | HK/CTRL | gata3 | ENSDARG00000016526 |
| lcp1 | HK/CTRL | lcp1 | ENSDARG00000023188 |
| hbba1 | HK/CTRL | ba1 | ENSDARG00000097238 |
| rag1 | HK/CTRL | rag1 | ENSDARG00000052122 |
| rag2 | HK/CTRL | rag2 | ENSDARG00000052121 |
| myca | HK/CTRL | myca | ENSDARG00000045695 |

| | | | |
|---|---|---|---|
| mycb | HK/CTRL | mycb | ENSDARG00000007241 |
| hmyc | HK/CTRL | hMYC | N/A |
| gfp | HK/CTRL | GFP | N/A |
| eef1a1l1 | HK/CTRL | ef1a | ENSDARG00000020850 |
| nkl.1 | NK cell | nkl.2 | ENSDARG00000058673 |
| nkl.3 | NK cell | nkl.3 | ENSDARG00000089202 |
| nkl.4 | NK cell | nkl.4 | ENSDARG00000044023 |
| nitr2a | NK cell | nitr2a | ENSDARG00000043398 |
| nitr2b | NK cell | nitr2b | ENSDARG00000000460 |
| nitr1b | NK cell | nitr1b | ENSDARG00000069848 |
| nitr1k | NK cell | nitr1k | ENSDARG00000110393 |
| nitr7b | NK cell | nitr7b | ENSDARG00000067977 |
| dicp1.1 | NK cell | dicp1.1 | ENSDARG00000091993 |
| ifng1-1 | ILC | ifng1-1 | ENSDARG00000045671 |
| ifng1-2 | ILC | ifng1-2 | ENSDARG00000024211 |
| il4 | ILC | il4 | ENSDARG00000087909 |
| il13 | ILC | il13 | ENSDARG00000077809 |
| il17a/f3 | ILC | il17a/f3 | ENSDARG00000041976 |
| il22 | ILC | il22 | ENSDARG00000045673 |

#### Supplementary Figure Legends

**Supplementary Figure 1: Lymphocytes in zebrafish scales. (A)** Scale light microscopy (top row, left panel for each genotype) and fluorescent microscopy (top row, right panel) images, from WT WIK and transgenic *cd79a:GFP*, *cd79b:GFP*, and *lck:eGFP* fish. Yellow boxes in top row correspond to high-power confocal images below. Confocal WT scale image shows an atypical ~30 µm faintly-fluorescent cell, presumably auto-fluorescence. White bars = 100µm in top row, 10µm in bottom row. **(B)** 3-dimensional confocal microscopy videos of scales from *cd79a:GFP*, *cd79b:GFP*, and *lck:eGFP* fish. *cd79a:GFP* and *cd79b:GFP* scale videos rotate left-to-right; *lck:eGFP* video presents z-stacks progressing from internal/medial (facing the body) to external/lateral (facing the water).

**Supplementary Figure 2: Gene expression profiles of lymphocytes from *cd79*-labeled lines. (A)** Copies of select **Figure 3A** heatmaps and corresponding dot-plots with different 2-gene indices, as indicated in the main text. **(B)** Transcript levels in FSC/SSC-based lymphoid gate GFP$^{-}$ thymic, scale, and marrow cells from *cd79a:GFP* and *cd79b:GFP* fish by multiplex-qPCR. Labels atop heatmaps indicate genotypes. *ef1a* was detected in all cells, indicating amplifiable RNA. Many, but not every (rightmost cells on each heatmap), cell expressed at least some lymphoid genes.

**Supplementary Figure 3: Gene expression profiles of P2 thymocytes of *cd79a:GFP* + *lck:mCherry* fish and GFP$^{hi}$ thymocytes of *lck:GFP* fish. (A)** Copy of **Figure 4B** thymus P2 heatmap, with dot-plot for *cd4* vs. *cd8*. **(B)** Copy of **Figure 4D** thymus GFP$^{hi}$ cell heatmap, with dot-plot for *cd4* vs. *cd8*. DP (pink; ~44%) and *cd8*$^{+}$SP (orange; ~41%) thymocytes are most abundant. **(C)** Transcript levels in GFP$^{lo}$ scale cells of *lck:GFP* + *rag2:hMYC* fish by multiplex-qPCR.

**Supplementary Figure 4: scRNAseq GEP of *lck:GFP* lymphocytes. (A)** Copy of UMAP plot of 1,890 $GFP^{hi}$ thymocytes and $GFP^{lo}$ scale and marrow lymphocytes from **Figure 6A**. **(B)** Heatmap of scaled expression of individual B-, T- and NK-lineage and housekeeping/control (HK/CTRL) genes, including leukocyte- (*lcp1*) and lymphoblast- (*rag*) specific transcripts and control [*GFP*, *myca*, *mycb*, *eef1a1l1* (labeled *ef1a*)] genes. ILC-specific genes (*ifng1-1*, *ifng1-2*, *il4*, *il13*, *il17a/f3*, *il22*, etc.) were near-absent in every population and are not shown. Select genes in Supplemental Table 2 were filtered out due to low expression in the entire data set. **(C)** Tissue-of-origin for each UMAP cluster. **(D)** UMAP plots divided by tissue-of-origin and color-coded based on collective lineage scores. **(E)** UMAP plots with S- and $G_2$/M-phase cell-cycle scores.

**Supplementary Figure 5: Scale biopsy does not substantially alter scale lymphocyte numbers.** Untreated **(A)** *cd79a:GFP*, **(B)** *cd79b:GFP*, and **(C)** *lck:GFP* fish had 5-scale biopsies, on opposite sides, 10 days apart. Upper panels show fluorescence microscopy fish images, with ImageJ fluorescence quantifications at right. Lower panels depict flow cytometry plots at left, with the number of lymphocytes recovered at right. White bars = 2 mm. Gates, cell quantities, and percent of total cells are shown in flow plots. n = number of scales analyzed.

**Supplementary Figure 1**

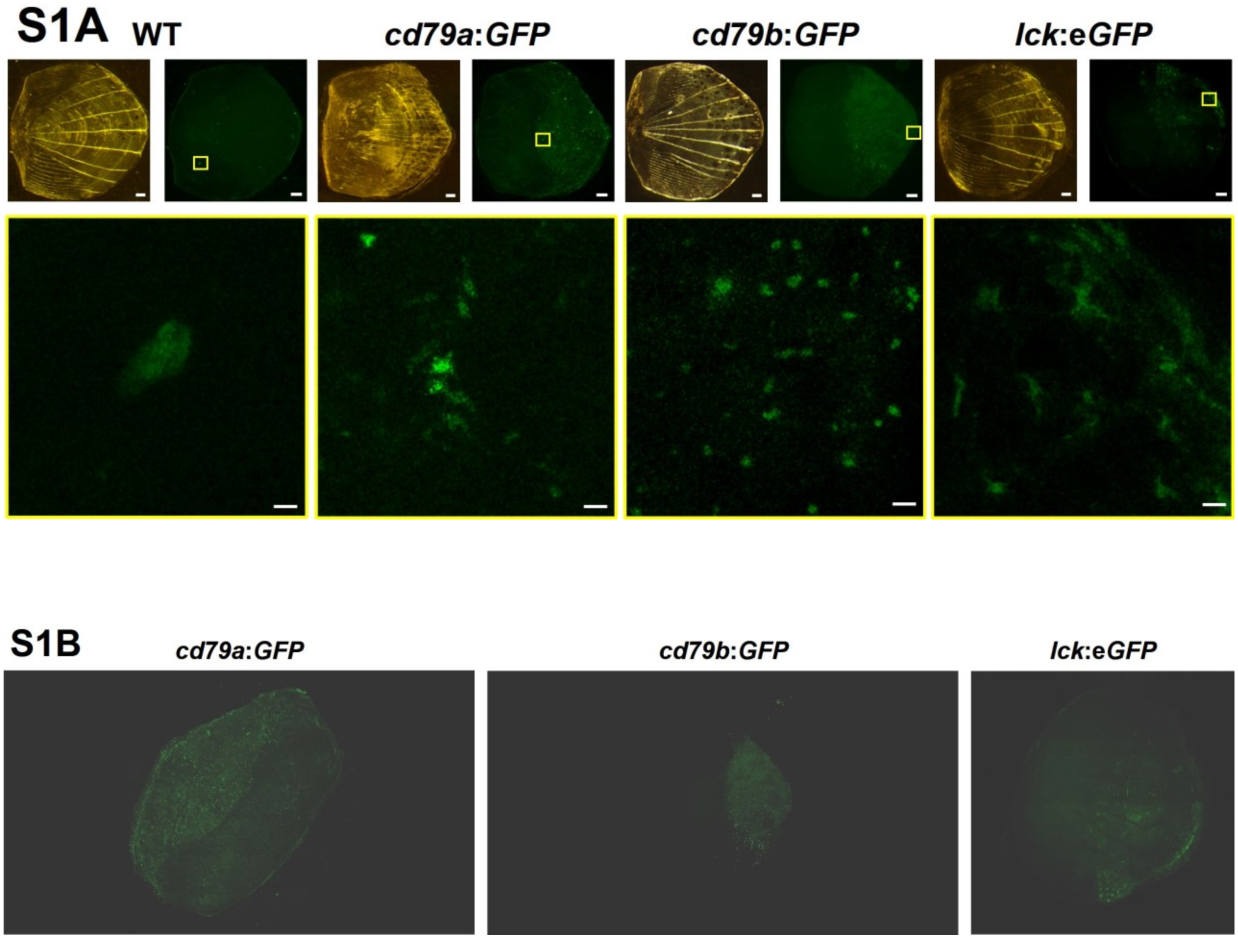

*cd79a:GFP* link: Check attached supplemental video file: “WT cd79a Scale#2 20x”

*cd79b:GFP* link: : Check attached supplemental video file: “WT cd79b scale#4 20x”

*lck:eGFP* link: Check attached supplemental video file: “WT lckGFP scale#1 20x”

### Supplementary Figure 2

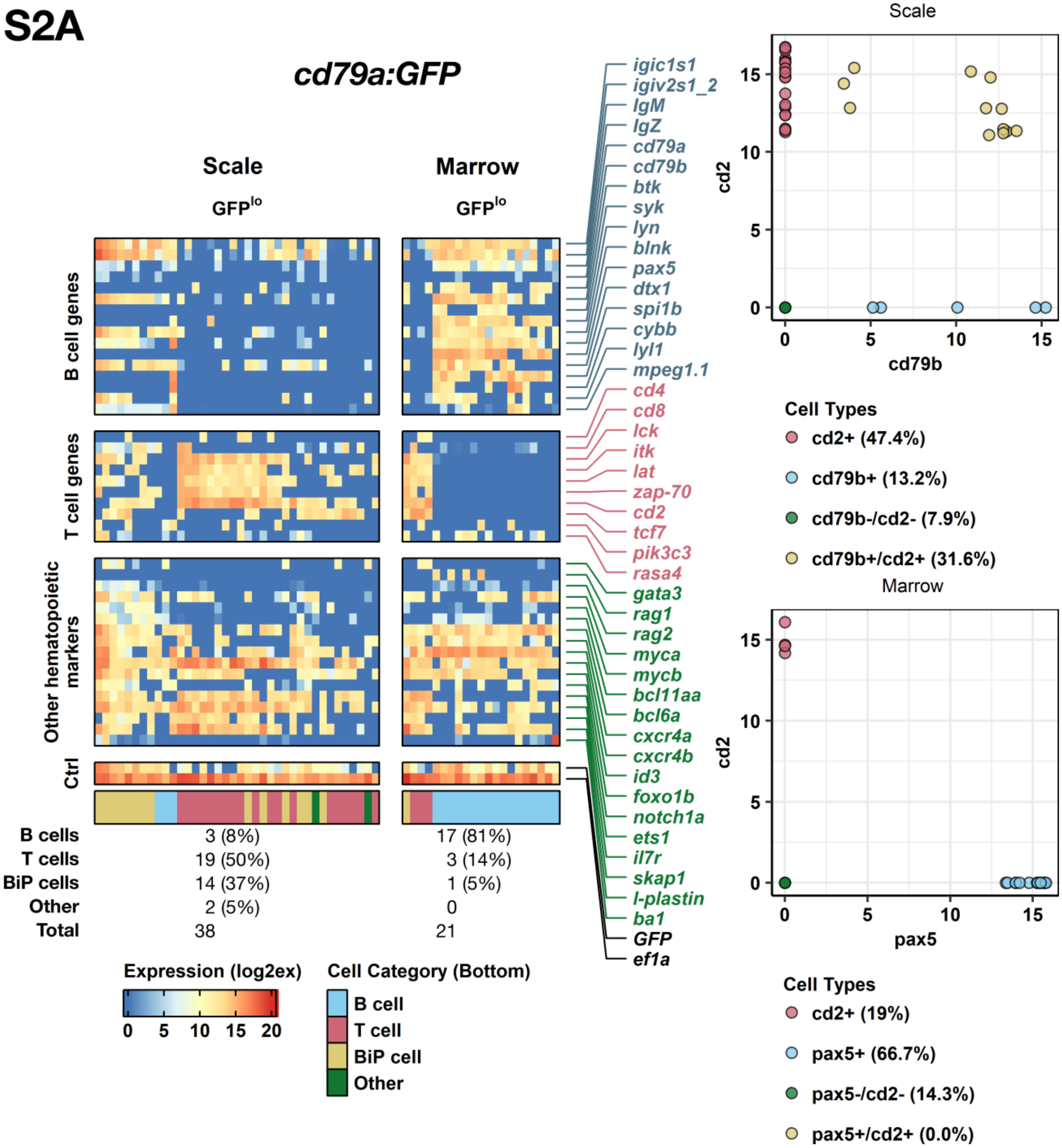

**S2B**

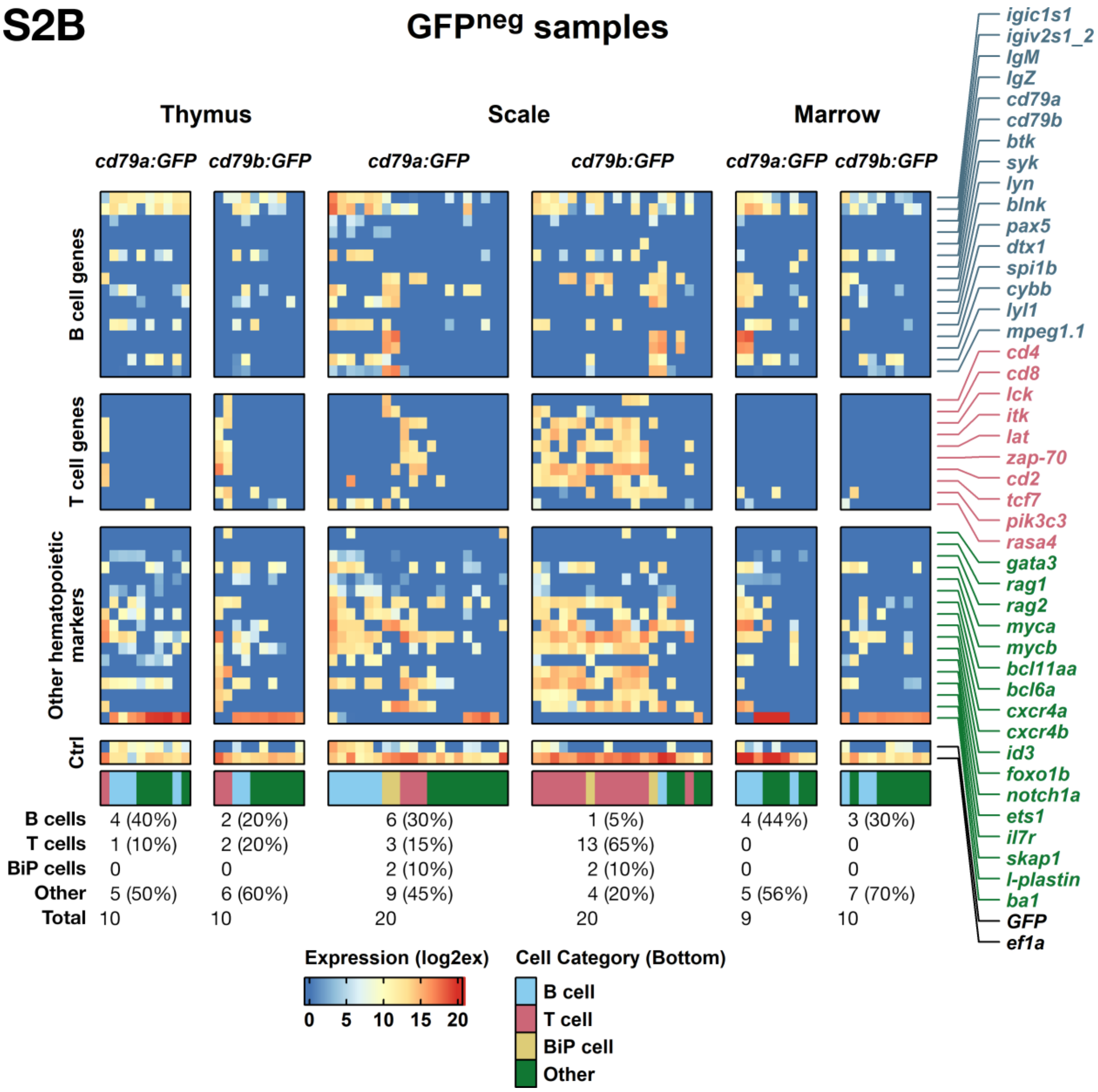
GFPneg samples
Thymus
Scale
Marrow
cd79a:GFP
cd79b:GFP
cd79a:GFP
cd79b:GFP
cd79a:GFP
cd79b:GFP
B cell genes
T cell genes
Other hematopoietic markers
Ctrl
igic1s1
igiv2s1_2
IgM
IgZ
cd79a
cd79b
btk
syk
lyn
blnk
pax5
dtx1
spi1b
cybb
lyl1
mpeg1.1
cd4
cd8
lck
itk
lat
zap-70
cd2
tcf7
pik3c3
rasa4
gata3
rag1
rag2
myca
mycb
bcl11aa
bcl6a
cxcr4a
cxcr4b
id3
foxo1b
notch1a
ets1
il7r
skap1
l-plastin
ba1
GFP
ef1a
B cells 4 (40%) 2 (20%) 6 (30%) 1 (5%) 4 (44%) 3 (30%)
T cells 1 (10%) 2 (20%) 3 (15%) 13 (65%) 0 0
BiP cells 0 0 2 (10%) 2 (10%) 0 0
Other 5 (50%) 6 (60%) 9 (45%) 4 (20%) 5 (56%) 7 (70%)
Total 10 10 20 20 9 10
Expression (log2ex)
0 5 10 15 20
Cell Category (Bottom)
B cell
T cell
BiP cell
Other

#### Supplementary Figure 3

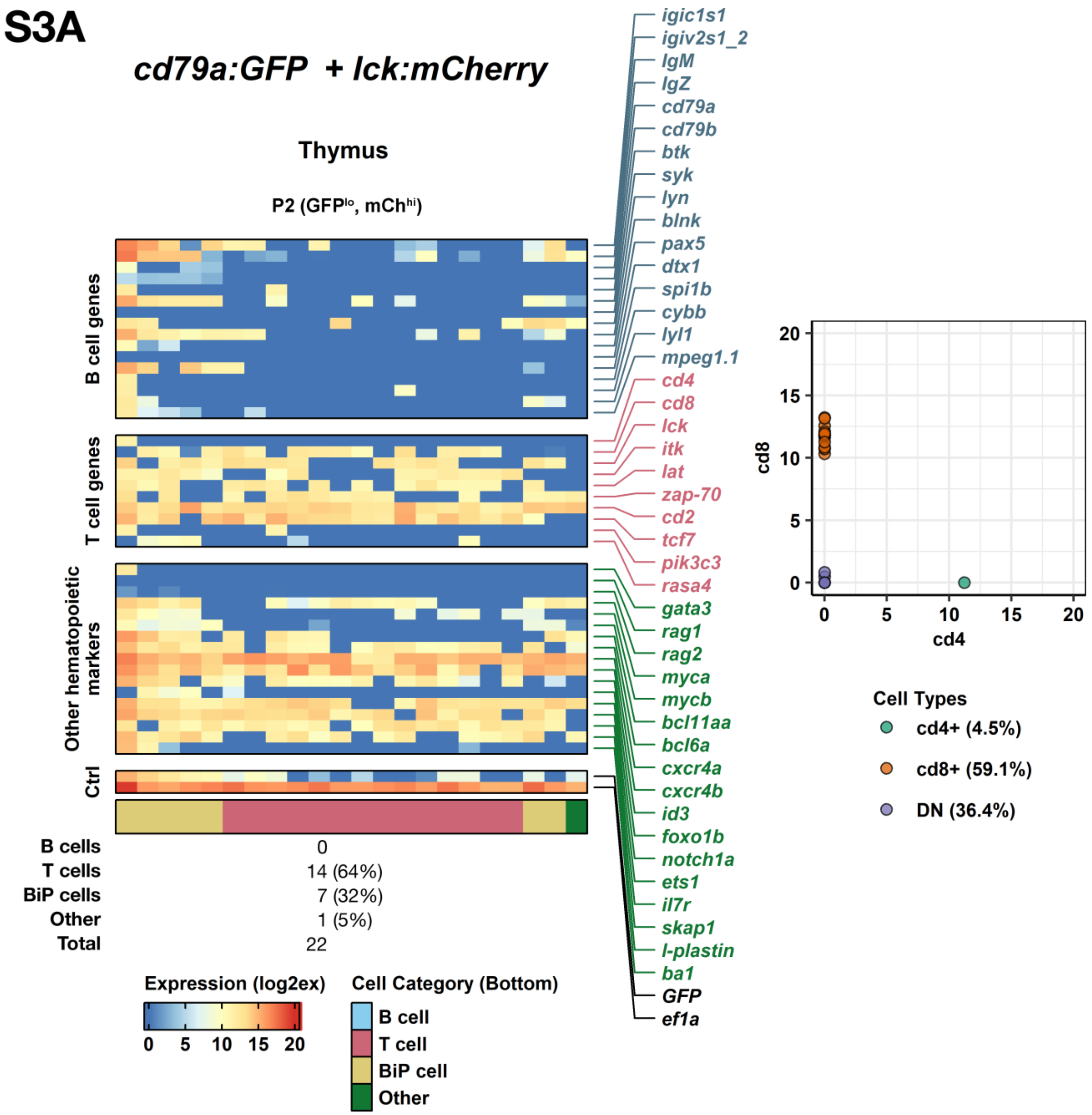

# S3B

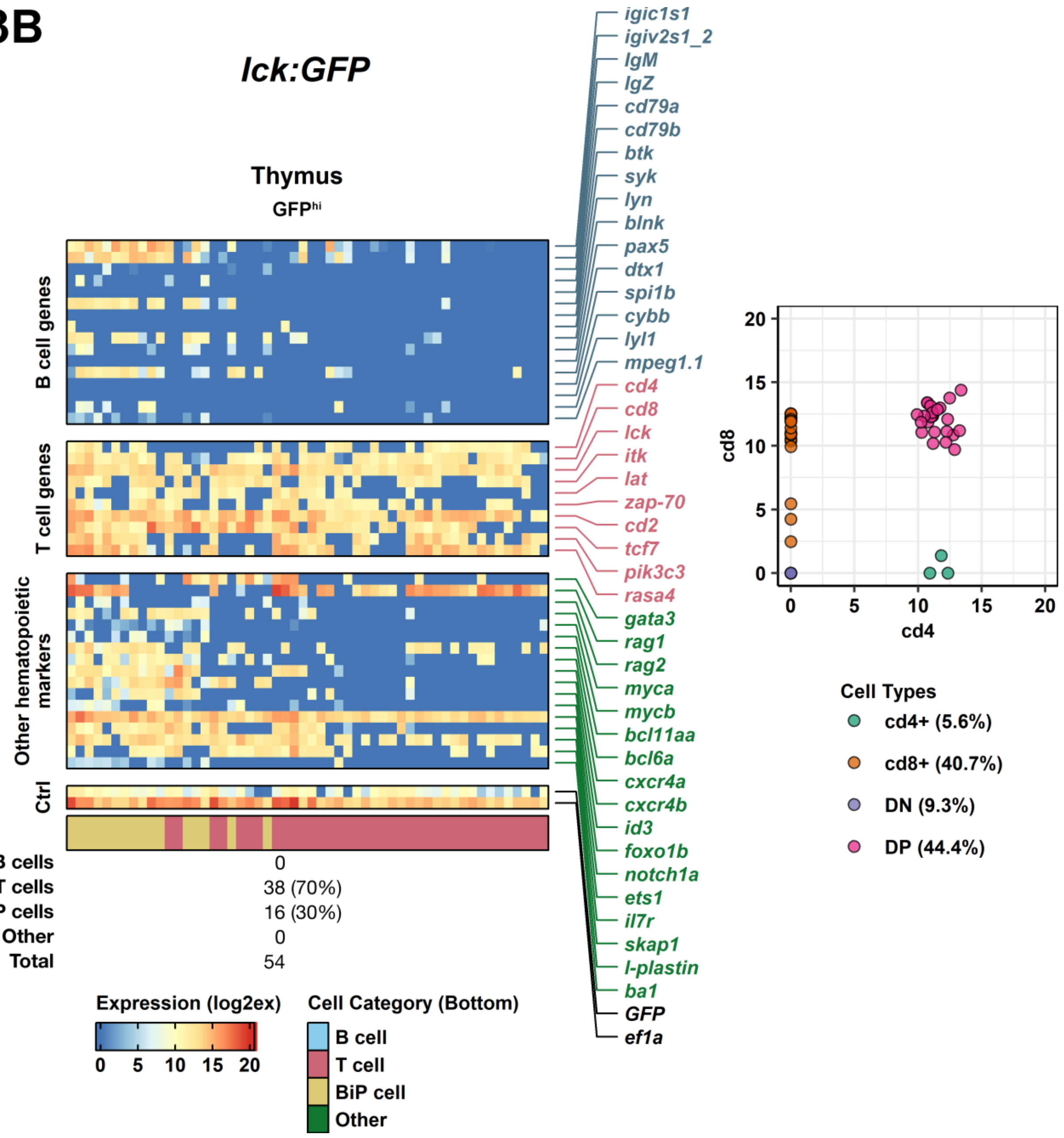
lck:GFP
Thymus
GFPhi
B cell genes
T cell genes
Other hematopoietic markers
Ctrl
igic1s1
igiv2s1_2
IgM
IgZ
cd79a
cd79b
btk
syk
lyn
blnk
pax5
dtx1
spi1b
cybb
lyl1
mpeg1.1
cd4
cd8
lck
itk
lat
zap-70
cd2
tcf7
pik3c3
rasa4
gata3
rag1
rag2
myca
mycb
bcl11aa
bcl6a
cxcr4a
cxcr4b
id3
foxo1b
notch1a
ets1
il7r
skap1
l-plastin
ba1
GFP
ef1a
B cells 0
T cells 38 (70%)
BiP cells 16 (30%)
Other 0
Total 54
Expression (log2ex)
0 5 10 15 20
Cell Category (Bottom)
B cell
T cell
BiP cell
Other
cd8
cd4
0 5 10 15 20
Cell Types
cd4+ (5.6%)
cd8+ (40.7%)
DN (9.3%)
DP (44.4%)

**S3C** ***lck:GFP + rag2:hMYC***

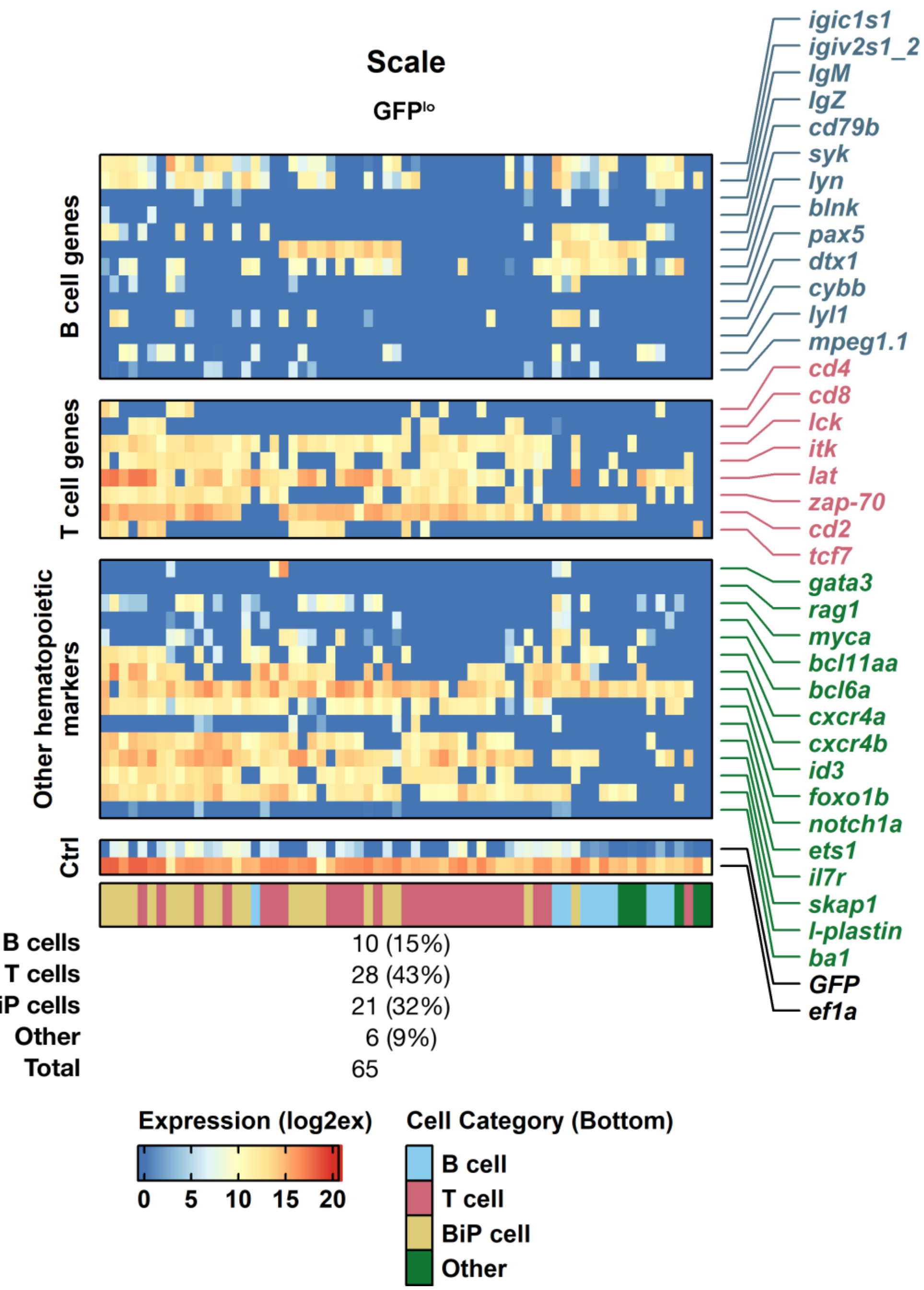

| | |
|---|---|
| **B cells** | 10 (15%) |
| **T cells** | 28 (43%) |
| **BiP cells** | 21 (32%) |
| **Other** | 6 (9%) |
| **Total** | 65 |

**Supplementary Figure 4**

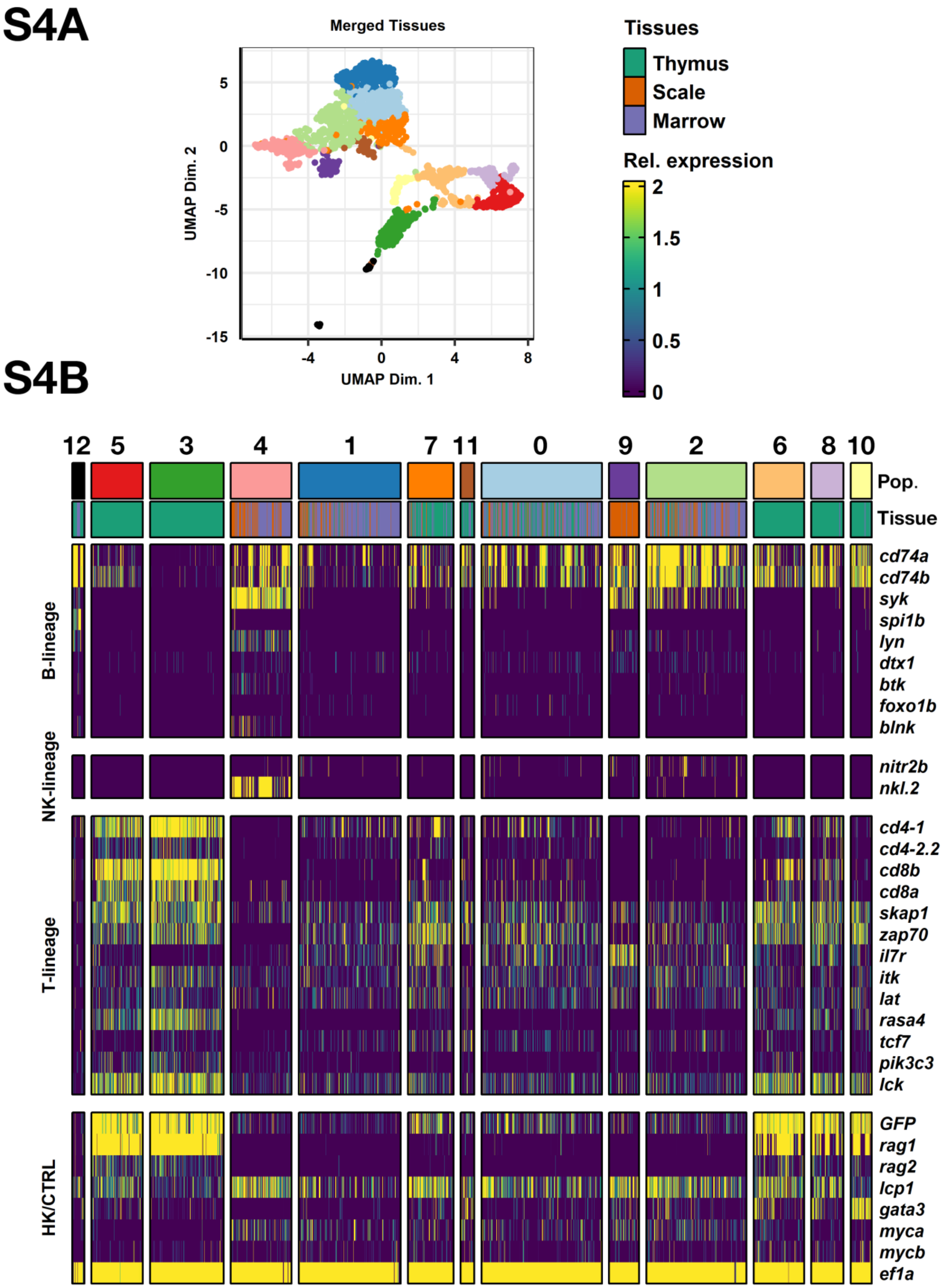

## S4C

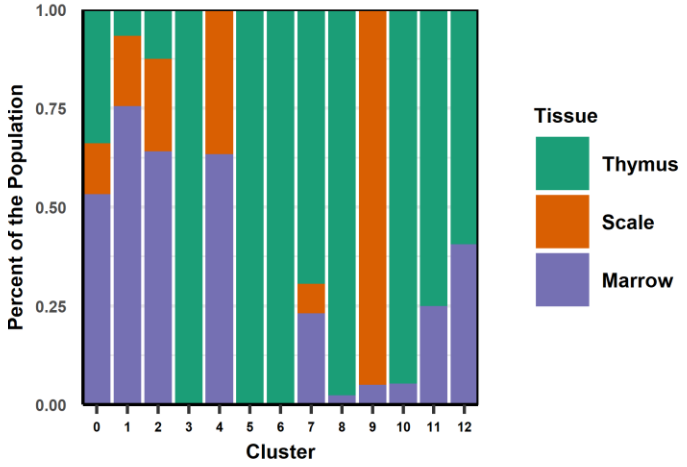

Percent of the Population
1.00
0.75
0.50
0.25
0.00
0 1 2 3 4 5 6 7 8 9 10 11 12
Cluster
Tissue
Thymus
Scale
Marrow

## S4D

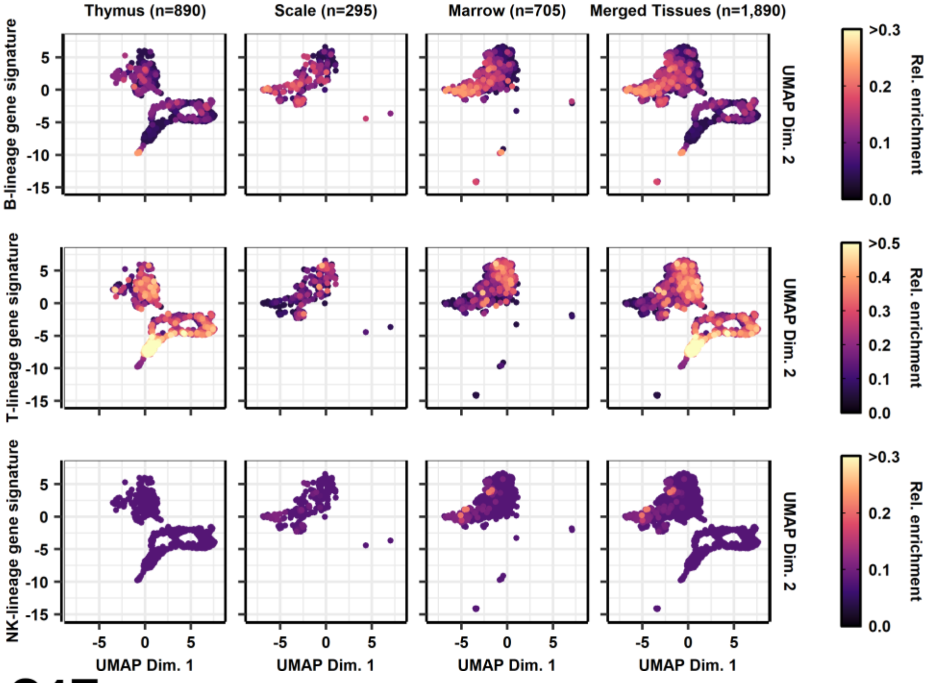

Thymus (n=890)
Scale (n=295)
Marrow (n=705)
Merged Tissues (n=1,890)
B-lineage gene signature
T-lineage gene signature
NK-lineage gene signature
UMAP Dim. 1
UMAP Dim. 2
Rel. enrichment
>0.3
>0.5

## S4E

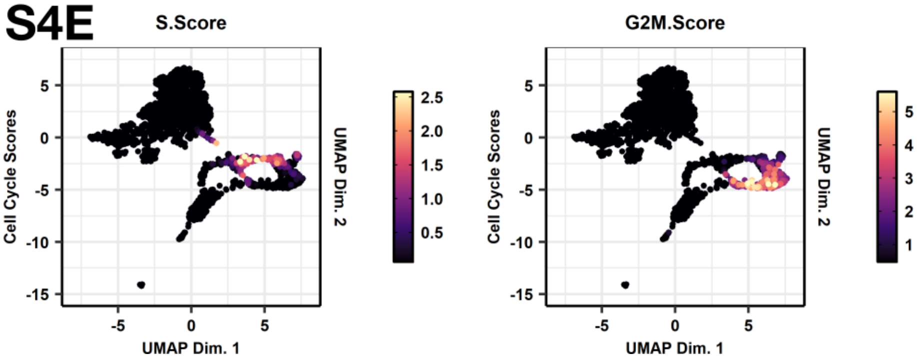

S.Score
G2M.Score
Cell Cycle Scores
UMAP Dim. 1
UMAP Dim. 2

### Supplementary Figure 5

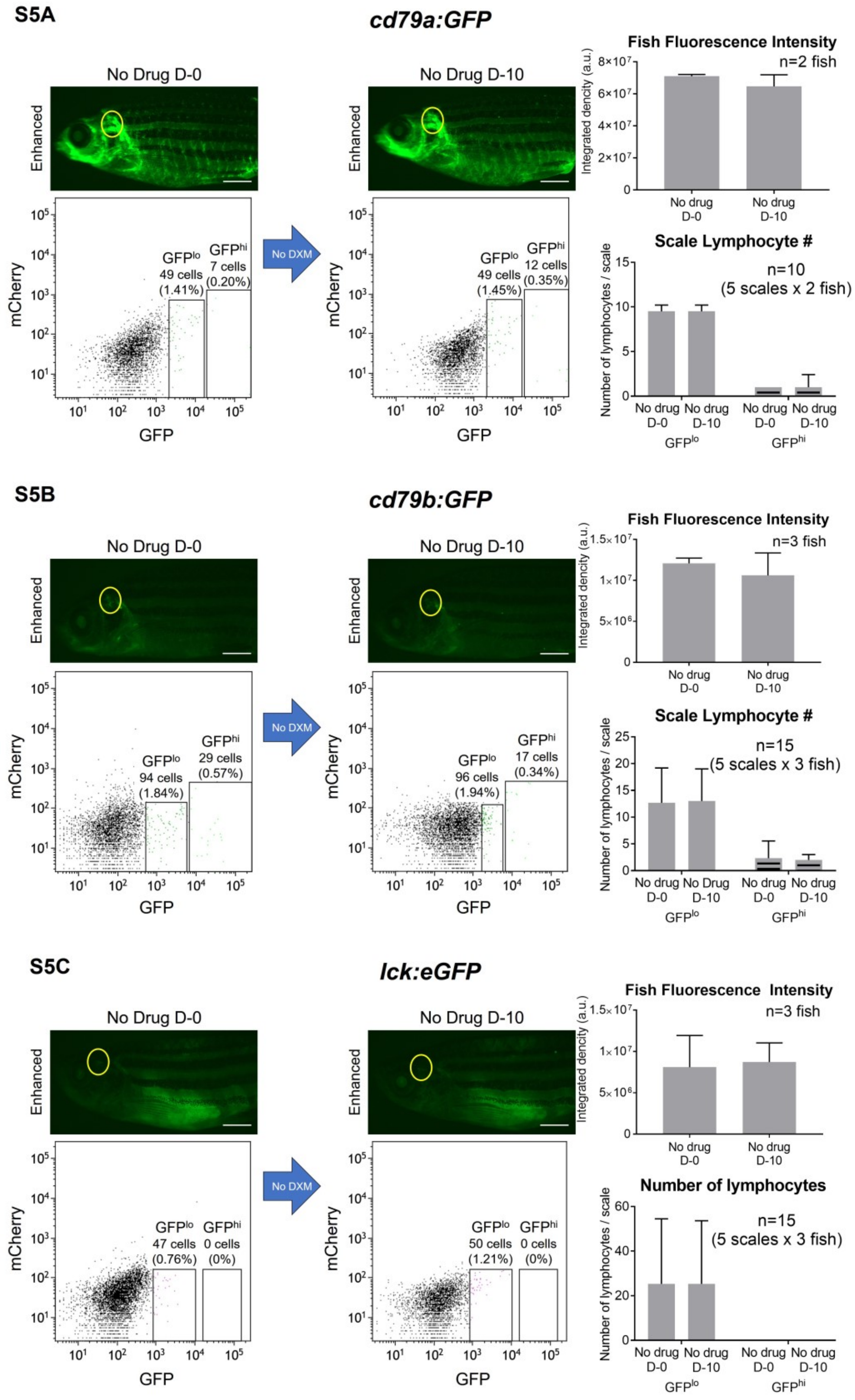

#### References:

1. Zheng GX, Terry JM, Belgrader P, et al. Massively parallel digital transcriptional profiling of single cells. *Nat Commun*. Jan 16 2017;8:14049. doi:10.1038/ncomms14049
2. Young MD, Behjati S. SoupX removes ambient RNA contamination from droplet-based single-cell RNA sequencing data. *Gigascience*. Dec 26 2020;9(12)doi:10.1093/gigascience/giaa151
3. Germain PL, Lun A, Garcia Meixide C, Macnair W, Robinson MD. Doublet identification in single-cell sequencing data using scDblFinder. *F1000Res*. 2021;10:979. doi:10.12688/f1000research.73600.2
4. Haghverdi L, Lun ATL, Morgan MD, Marioni JC. Batch effects in single-cell RNA-sequencing data are corrected by matching mutual nearest neighbors. *Nat Biotechnol*. Jun 2018;36(5):421-427. doi:10.1038/nbt.4091
5. Zappia L, Oshlack A. Clustering trees: a visualization for evaluating clusterings at multiple resolutions. *Gigascience*. Jul 1 2018;7(7)doi:10.1093/gigascience/giy083
6. Hao Y, Hao S, Andersen-Nissen E, et al. Integrated analysis of multimodal single-cell data. *Cell*. Jun 24 2021;184(13):3573-3587 e29. doi:10.1016/j.cell.2021.04.048
7. Andreatta M, Carmona SJ. UCell: Robust and scalable single-cell gene signature scoring. *Comput Struct Biotechnol J*. 2021;19:3796-3798. doi:10.1016/j.csbj.2021.06.043
8. Wu T, Hu E, Xu S, et al. clusterProfiler 4.0: A universal enrichment tool for interpreting omics data. *Innovation (Camb)*. Aug 28 2021;2(3):100141. doi:10.1016/j.xinn.2021.100141
9. Liberzon A, Subramanian A, Pinchback R, Thorvaldsdottir H, Tamayo P, Mesirov JP. Molecular signatures database (MSigDB) 3.0. *Bioinformatics*. Jun 15 2011;27(12):1739-40. doi:10.1093/bioinformatics/btr260
10. Lopez D, Montoya D, Ambrose M, et al. SaVanT: a web-based tool for the sample-level visualization of molecular signatures in gene expression profiles. *BMC Genomics*. Oct 25 2017;18(1):824. doi:10.1186/s12864-017-4167-7
11. Hu C, Li T, Xu Y, et al. CellMarker 2.0: an updated database of manually curated cell markers in human/mouse and web tools based on scRNA-seq data. *Nucleic Acids Res*. Jan 6 2023;51(D1):D870-D876. doi:10.1093/nar/gkac947
